# Persistent effects of dietary selection and inbreeding on microbiome composition and longevity in Drosophila

**DOI:** 10.1101/2025.04.26.650496

**Authors:** Peyton K. Warren, Garrison Miller, Gaurav S. Kandlikar, Enoch Ng’oma

**Affiliations:** Division of Biological Sciences, University of Missouri-Columbia, MO, USA; Department of Biological Sciences, Louisiana State University, LA

**Keywords:** experimental evolution, nutritional history, longevity, microbiome legacies, dietary stress, persistent effects

## Abstract

**Background:** Dietary environments can shape host-microbe co-evolution by imposing selection pressures that favor host and microbial genotypes with enhanced fitness under specific nutritional conditions. However, the long-term evolutionary dynamics of these interactions after selection pressures are relaxed remain poorly understood. Here, we investigated the microbiome of *Drosophila melanogaster* populations that underwent long-term experimental evolution under three temporally variable nutritional regimes (constant high sugar, progressively decreasing protein, and fluctuating protein), and compared them to unselected controls. Following inbreeding of evolved populations, we examined how persistent evolutionary effects interact with aging to shape host-microbe relationships and survival.

**Results:** All selected regimes exhibited reduced survival relative to controls, indicating lasting physiological costs associated with historical selection and inbreeding. Survival differed among regimes: the deteriorating-protein regime was closest to controls, the fluctuating-protein regime was intermediate, and the high-sugar regime showed the shortest lifespan. Survival effects were sex-specific: relative to control females, those from the fluctuating-protein regime exhibited reduced early-life survival, whereas females from the deteriorating-protein regime experienced greater late-life mortality. Microbiome composition varied with both selection regime and age. Although Firmicutes and Proteobacteria dominated across groups, selected lines showed increased Firmicutes and reduced Proteobacteria, especially early in life, suggesting early-life taxonomic restructuring. The decreasing-protein regime maintained more stable microbial diversity over time, whereas high-sugar and fluctuating-protein diets were associated with progressive microbiome instability with age. Core *Acetobacter* species (*A. aceti*, *A. oryzifermentans*) declined in abundance in selected flies, indicating persistent disruption of microbiome integrity. A random forest model predicted fly age from microbiome composition with 78.8% accuracy, reinforcing links between microbial dynamics and host aging.

**Conclusion:** Historical dietary selection and inbreeding can leave lasting signatures on survival and microbiome composition in *Drosophila*. Protein restriction promoted late-life longevity and microbial stability, whereas high-sugar and fluctuating diets were associated with early-life effects followed by later-life shifts in dominant taxa. Together, these findings illustrate how nutritional history and its microbial legacies influence lifespan and aging through persistent host-microbe interactions.

## Introduction

Microorganisms are deeply integrated into host biology across taxonomic and functional scales, playing critical roles in organismal performance and fitness (Ley et al. 2008; Muegge et al. 2011; Gould et al. 2018). Rapidly changing environmental conditions increase the pressure for hosts to cope with novel conditions faster than their genomes can evolve. Microbial genomes, which evolve more rapidly than host genomes, represent an important source of ecological and functional flexibility, enabling hosts to respond to environmental stressors through shifts in symbiotic composition and activity. Microbiotas, as part of the functional hologenome, can mediate developmental timing, immune defense, energy allocation, and other traits central to host survival and reproduction [4, 5]. In many species, the microbiome is a key modulator of host fitness [6], development [7, 8], reproduction [9, 10], stress tolerance [11], pathogenic defense [12, 13], survival, and aging [13–15]. Disruptions to microbial composition can increase gut permeability and systemic inflammation, accelerating physiological decline [16], whereas beneficial microbes can attenuate immune overactivation and extend lifespan [14]. These findings position inflammation as a central axis through which microbiotas mediate life-history trade-offs, particularly under dietary stress [14, 16], reviewed in [17, 18].

Diet is a dominant ecological determinant of host-associated microbiomes. In *Drosophila melanogaster*, while variation in microbiome composition has been linked to host allele frequency shifts in genes associated with local adaptation [4], microbial composition is often more strongly influenced by the nutritional environment than host genotype [19, 20]. For example, *Lactobacillus plantarum* enhances larval growth on poor diets by promoting insulin signaling [7], while acetic acid bacteria regulate sugar metabolism and immune function [21]. Nutrient availability thus shapes both microbiota composition and downstream host traits including immune investment, somatic maintenance, and fecundity. Many studies have shown that reduced protein-to-carbohydrate (P:C) ratios extend lifespan at the cost of reproductive output, whereas protein-rich diets promote early-life fecundity but reduce survival [22–24]. These physiological responses are mediated through nutrient-sensing pathways such as IIS and mTOR [25, 26], which are also modulated by gut microbes [27]. Given this complex interplay, sustained dietary stress is expected to shape host physiology and microbiome composition in concert.

Intrinsic host factors such as age also interact with microbial dynamics. In *D. melanogaster*, aging is associated with shifts in gut microbiota composition, including reduced diversity and expansion of inflammatory taxa [14, 28]. These changes are not simply passive consequences of aging but actively contribute to age-related physiological decline, suggesting that the microbiome is both a target and mediator of aging processes.

Here, we examine a set of experimentally evolved *D. melanogaster* populations that were maintained for 65 generations under distinct dietary selection regimes and subsequently inbred for >30 generations due to COVID pandemic-related disruptions. This design enables us to investigate whether long-term dietary selection has left lasting genomic or microbiome signatures that persist after extended inbreeding and rearing under a common diet. Selection treatments included: 1) deteriorating availability (DA), representing high-to-low protein provision within each generation; 2) fluctuating availability (FA), reflecting predictable high-low-high protein cycles; and 3) constant high sugar (CH), modeling persistent metabolic excess. A control group (C) was maintained without dietary stress.

Our primary goal was to determine whether historical dietary selection is reflected in contemporary microbiome composition and host survival patterns. We therefore refer to any observed differences as *legacies of dietary history* rather than direct evidence of ongoing adaptation. We hypothesized that hosts derived from contrasting nutritional histories would retain distinct microbiome compositions reflecting their ancestral environments, potentially influencing lifespan through pathways linked to inflammation and metabolic homeostasis.

Specifically, we predicted that the DA regime would show signatures consistent with investment in somatic maintenance and longevity, potentially mediated by microbes that mitigate inflammatory processes. In the FA regime, evolved under predictable cycles of abundance and scarcity, we anticipated intermediate survival outcomes and dynamically shifting microbial compositions reflecting metabolic flexibility. Chronic high-sugar exposure (CH regime) was expected to produce signatures of metabolic stress and reduced microbial diversity. Finally, we anticipated that age-related microbiota succession would interact with these dietary legacies, producing distinct temporal patterns of microbial resilience or instability.

Collectively, these predictions provide a framework for testing how long-term nutritional history, filtered through both host and microbial evolution, can leave enduring effects on host–microbiome interactions and life-history dynamics, even after prolonged inbreeding and common-garden rearing.

## Materials and methods

### Experimental evolution

As part of a larger study, we experimentally evolved a population of *D. melanogaster* under several stressful nutritional conditions. We generated the initial population by mass-intercrossing 835 recombinant inbred lines from the B population (pB) of the *Drosophila* Synthetic Population Resource, DSPR [29, 30] for five generations under common garden conditions (Fig. 1a) [31]. Several Drosophila community panels in North America, including *Drosophila* Genetic Reference Panel (DGRP) and the *D. melanogaster* Reference Genome, exhibit mosaic genomic ancestry (∼20% African, ∼80% European) shaped by natural selection against incompatible allele combinations [32]. This can also be expected in DSPR populations, highlighting the need for a genetically diverse population in experimental evolution, as such expanded genotypic diversity would provide a broad substrate for selection facilitating the study of complex traits and adaptation. The properties of the DSPR are described elsewhere [29, 30]. From this genetically diverse base population, 36 lines were independently subjected to one of three dietary selection regimes for 65 generations with each regime replicated 12 times in large population cages (N > 4,000 flies; Fig. 1a). One population (N > 4,000) was maintained without selection exposure for the duration of selection. Selection in these populations acted primarily on reproductive success and early-life survival under dietary stress, rather than directly on lifespan. Consequently, differences in longevity observed in the inbred lines likely reflect correlated responses to selection on nutrient-dependent fitness rather than direct selection for extended lifespan.

**Figure 1:**
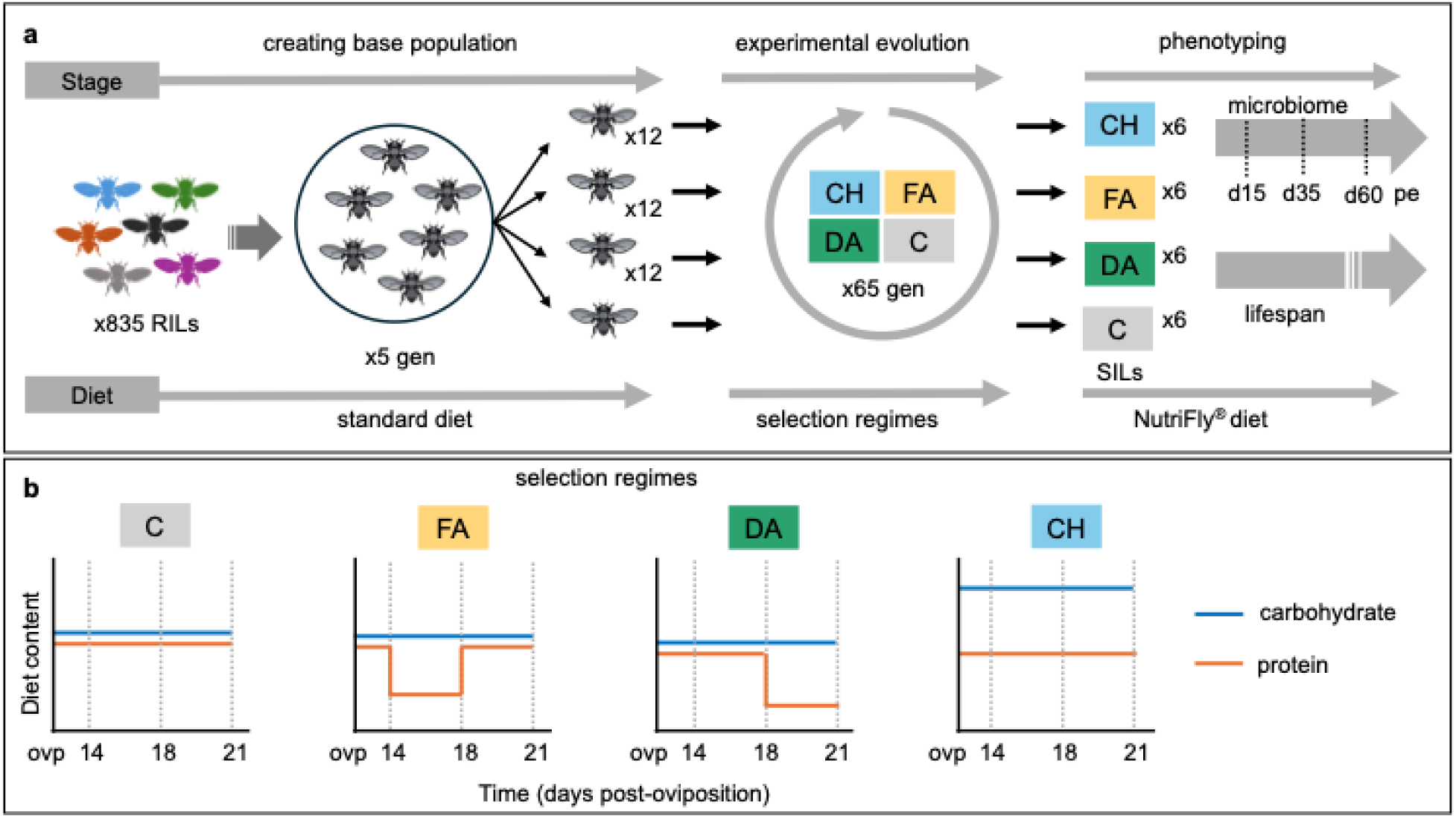
Study design. **a.** A total of 835 recombinant inbred lines of the DSPR rared on standard maintenance diet were interbred into a genetically diverse population and subjected to experimental evolution in four diet treatments for 65 generations. Selection was terminated after 65 generations, and flies in each regime inbred in vial replicates, here denoted, *selection inbred lines*, or SILs and maintained on common NutriFly diet. Control flies were maintained on standard diet throughout selection in one cage at N > 4,000 and also moved to NutriFly at selection termination. Six SILs per regime were each expanded into 6 replicates, alongside pseudo replicate vials of C flies. Microbiome composition was measured at 15, 35 and 60 days post-eclosion in 3 vial replicates. Survival was measured in a separate set of 3 vials. **b.** Selection regimes: 1) fluctuating (FA), 2) deteriorating (DA) and 3) constant high (CH) nutrient availability were set up in 12 replicate cages (N > 4000 flies per cage) and fed sequences of low protein (LP) and control (C) diet alternations or constant high sugar (HS) diet over a 21-day cycle post-oviposition (ovp) per generation (see methods). Note: DA and FA received temporal alternations of contro and low protein levels; CH received an oversupply of sugar; C was retained on standard diet for the DSPR (no deliberate selection).

#### Nutritional composition of diets constituting selection regimes

Based on previously published formulations [25, 31, 33], we used three diets that systematically varied in the ratio of protein (yeast) to carbohydrate (sucrose) to impose distinct nutritional selection pressures. Each diet was prepared per liter of water with the following composition: Control diet - 10 g agar, 50 g sucrose, 200 g yeast, and 2.7 g tegosept (dissolved in 11 mL ethanol); Low-protein diet - 10 g agar, 50 g sucrose, 100 g yeast, and 2.7 g tegosept; High-sugar diet - 10 g agar, 342 g sucrose, 200 g yeast, and 2.7 g tegosept. Yeast provided the principal protein source, and sucrose supplied carbohydrates. The control diet yielded an intermediate protein: carbohydrate ratio (P:C ≈ 1: 0.5) typical of standard laboratory diets that support optimal growth and fecundity.

The low-protein diet reduced yeast content by half while maintaining sucrose levels, generating a protein-limited condition (P:C ≈ 1: 1). The high-sugar diet increased sucrose concentration roughly sevenfold, creating a carbohydrate-rich environment (P:C ≈ 1: 3). Percent macronutrient content by energy was approximately 45 - 53% protein and 47–52% carbohydrate for the control, 36–41% protein and 59–64% carbohydrate for the low protein diet, and 17–19% protein and 81–83% carbohydrate for high sugar diet.

#### Maintenance of the control population

Prior to the onset of experimental evolution, the recombinant inbred lines used to generate the base population were maintained on a standard cornmeal–dextrose maintenance diet (40.8 g cornmeal, 21.6 g yeast, 86.3 g dextrose, 6.25 g agar per liter of water, with 1.8 g tegosept). This base diet contained lower protein and carbohydrate levels than any of the experimental diets, ensuring that the defined sugar–protein diets represented clear nutritional shifts.

To preserve an ancestral baseline for comparison, the unselected control population was maintained throughout the experiment on the cornmeal–dextrose maintenance diet described above - the same diet on which the DSPR founder population was reared prior to the start of selection. We note that this “maintenance” diet differs from the “control (C)” diet used as one of the experimental selection treatments (see above). Using the ancestral diet for the unselected population ensured that these flies represented the pre-selection reference state, avoiding inadvertent selection under any of the experimental diets. The control diet was considered optimal for growth and reproduction and provided a reference for evaluating nutritionally stressful regimes (low protein or high sugar) that impose either limitation or oversupply of key nutrients (previously characterized in Ng’oma et al. [34]).

#### Construction of selection treatment regimes

From these diets, we designed three selection conditions (Fig. 1b) by alternating the sequence in which the diets were presented to flies in each selection generation. The Fluctuating Availability (FA) treatment alternated nutrient provision as follows: flies received a control until day 14 post-oviposition (po), low protein diet from day 14 - 18 po, and control diet again until day 21 po when eggs were collected to seed the next generation. Flies in the Deteriorating Availability treatment (DA) received a standard diet until day 18 po, followed by a low protein diet until day 21 po, at which eggs were collected. Files in the Constant High sugar treatment (CH) received the high sugar diet throughout, until day 21 po. Alongside selection cages, we maintained one population (N > 4000) that was not exposed to experimental selection and likely remained outbred at the time of this assay, with no observed population bottlenecks and assuming weak genetic drift. Because eggs were collected from survivors at day 21 po each generation, selection acted primarily through differences in fecundity and viability during early adulthood under each dietary regime. Longevity beyond this window was therefore not a direct target of selection but could evolve indirectly as a correlated response to nutrient-dependent reproductive success and stress tolerance. These dietary regimes therefore varied in both macronutrient composition and temporal availability during selection. This may invoke distinct evolutionary responses in the reproduction–survival trade-off. We interpret results with this fact in mind.

#### Selection inbred lines (SILs)

Experimental evolution was performed for 65 generations, after which populations were inbred in replicated vials for each of 36 selection populations at the onset of the Covid pandemic for 38 generations (∼19 months). At the end of experimental evolution, all flies were of similar age and sampled from the actively reproducing cohort, ensuring that alleles influencing fecundity and early-life survival were those most consistently transmitted to the next generation. We refer to inbred lines derived from experimental selection as Selection Inbred Lines (SILs, Fig. 1a). SILs were maintained in common garden conditions on a molasses-based Nutri-Fly® diet (Genesee Scientific) until assaying, and the unselected group was held at high population in a cage until assay.

#### Sample collection

For this study, we selected six random SILs from each selection regime (FA, DA, and CH) along with 12 pseudo replicated vials of the non-selected population. Each selected SIL was expanded over three generations, and three vials each were assigned to microbiome and lifespan estimation assays. Each assay vial was seeded with 24 females and 6 males to reduce male mating pressure, which is known to reduce female lifespan in *Drosophila* [34, 35]. This set up minimizes mating-induced costs and allows clearer inference on intrinsic female lifespan differences among selection treatments. Our design therefore yields a reduced male sample in lifespan analysis which we consider in the interpretation of results (i.e., 1,670 deaths, 176 censored events for females; 572 deaths, 9 censored events for males). Death events were counted three times per week (Monday, Wednesday, Friday) at which time also a fresh food vial was provided. For the microbiome portion, we assayed only females. Based on our prior study indicating key variability in lifespan and fecundity responses to diet across life phases [36], we collected whole flies at three time points (15, 35, and 60 days po, Fig. 1). At each time point, four females were flash-frozen and stored at -80°C for analysis.

#### Diet sources

Diets were prepared with inactivated SAFPro Relax + YF 73050 yeast (Lesaffre Yeast Corp., Milwaukee, USA), stored at 4°C, and used within 14 days. Evolved lines were held on a common Nutri-fly molasses formulation (Genesee Scientific) based on a Bloomington Drosophila Stock Center Nutri-fly molasses protocol (12.4 g/L yeast, 61.3 g/L yellow cornmeal, and 75.2 ml/L molasses). This change was necessitated by shortages in supplies during and after the pandemic. Across all assays, flies were maintained in a temperature-controlled chamber at 23°C under continuous light, mirroring conditions under which the DSPR was originally generated and maintained [29]. Flies were supplied with fresh food three times per week (Monday, Wednesday, Friday).

### 16S rRNA amplicon sequencing

Whole-fly 16S rRNA sequencing without surface sterilization captures both internal and surface-associated microbes and thus represents the total host-associated microbiome. This approach aligns with our goal of characterizing microbiome responses to long-term dietary selection at the whole-organism level. All samples were reared on a common diet medium prepared in bulk and distributed across all selection lines, ensuring a shared environmental microbial background. Therefore, observed differences across selection treatments likely reflect evolved host-microbiome associations rather than environmental contamination. DNA was extracted from pools of two to four flies using the Qiagen DNeasy Blood & Tissue Kit (Cat. No 69504). Samples were manually homogenized with a glass pestle in ATL buffer, incubated for 4 hours, and processed following the kit manufacturer’s protocol. DNA yield and quality were assessed using a nanodrop spectrophotometer (ND-1000) and Qubit 4 Fluorometer with the Qubit 1X dsDNA HS Assay Kit (Cat. No Q33231). Samples were normalized as needed with elution buffer and sequenced on a MiSeq instrument (PE250-v2 Nano flow cell, 25 cycles) targeting the V4 region of the bacterial 16S rRNA gene with primers 515F (5’-GTGCCAGCMGCCGCGGTAA-3’) and 806R (5’-GGACTACHVGGGTWTCTAAT-3’) [37, 38]. Sequencing was performed at the University of Missouri Genomics Technology Core.

### Data Analysis

#### Lifespan

We first explored and compared survival patterns among selection regimes in a total of 2,427 individuals (2,242 deaths, 185 censored), using survival models that included sex either as a covariate or in sex-stratified analyses. Kaplan-Meier survival curves indicated marked sex differences in survival probability (Additional File 2 Table S1) and changes in the relative survival rankings of selection regimes over time. Because each selection regime (CH, DA, FA) consisted of multiple unique selection inbred lines (SILs) replicated across vials, our design required modeling fixed effects of regime, sex, and their interaction, while accounting for random effects of SIL and vial.

To evaluate the effects of dietary regime and sex on survival, we fitted mixed-effects Cox proportional hazards models with a random intercept for SIL nested within vial. We compared an additive model (`event ∼ regime + sex + (1 | SIL/vial)`) and an interaction model (`event ∼ regime * sex + (1 | SIL/vial)`) using ΔAIC as the comparison criterion. Proportional hazards assumptions were tested using Schoenfeld residuals. All fixed effects (regime, sex, and their interaction) significantly violated this assumption (χ² test, *p* < 0.01), indicating time-varying hazard ratios. A time-transformed Cox model (`regime + tt(sex)`, where `tt(x) = x * log(t)`) improved fit (ΔAIC = 15.6), confirming time-dependent effects of sex.

Given these violations and the hierarchical structure of our data, we proceeded with a discrete-time survival framework. We implemented a generalized linear mixed model (GLMM) with a complementary log–log (cloglog) link using **lme4**, specifying event ∼ regime * sex + (1 | SIL). This model structure best fit the data (ΔAIC = 17, *p* = 2.7 × 10⁻⁵), while additional vial-level random effects did not improve fit (ΔAIC = +2).

To confirm robustness, we re-estimated the same model using a fully Bayesian hierarchical approach implemented in **brms**, specifying identical fixed and random effects and a cloglog link. Weakly informative priors were applied (normal(0, 2) for fixed effects; Student-t(3, 0, 2.5) for random effects). Results from both approaches were concordant, and the Bayesian model was used for inference and visualization. Full model specifications and diagnostics are provided in Additional File 1.

Our recent study testing effects of larval nutrition on adult phenotypes in this population [36] showed that patterns of survival can vary significantly across shorter periods of time in adult lifespan. To understand evolved patterns of survival in each selection treatment, we examined percentile survival differences in ten-day intervals in which each of our sampling time points for the microbiome portion (described next) formed the median: 10-20, 30-40, and 55-65 days, post eclosion. We tested for significant differences between treatments in each time interval I1, I2 and I3 using either analysis of variance (Anova) or Kruskall-Wallis tests followed by an appropriate post-hoc test (Turkey’s HSD or Dunn’s Test).

#### Microbiome analysis

We processed Illumina reads using the R library DADA2 pipeline [39], leveraging its use of Amplicon Sequence Variants (ASV), an alternative to Operational Taxonomic Units (OTUs) that enables the detection of single-nucleotide differences in amplicon sequencing and avoids the limitations of traditional OTU clustering by using exact biological sequences. Reads were quality-filtered and trimmed at positions 225 (forward) and 200 (reverse), and chimeras were removed. Taxonomy was assigned using the Silva database (v138.1). Samples with fewer than 200 reads after sequence merging were excluded. We investigated the proportion of reads identified as *Wolbachia* and found that these constituted less than 1% reads in the entire data set (Additional File 3, Fig. S7). Prior to analysis, these reads were removed as common practice [40]. We performed mainstream analysis and visualization using the R library **phyloseq** [41]. Read counts were normalized using a log-transformation: `log(1 + x)/sum(log(1 + x))`. This approach combines two commonly used normalization equations, log(1 + x) and (x)/sum(x), into one step and effectively scales data to relative abundances (i.e., the proportion of specific microbial groups within samples) while revealing differences in rare taxa through its log transformation [42–44]. This method was used instead of rarefication because rarefication is known to unnecessarily discard samples and to have a higher false positive rate (McMurdie and Holmes 2014).

We assessed microbiome diversity using both alpha and beta diversity metrics. For alpha diversity, we calculated Shannon and observed species indices from non-normalized sequence data, as standard practice to preserve true richness and evenness without introducing artifacts from normalization. Beta diversity was also evaluated on a full data set using principal coordinate analysis (PCoA) based on UniFrac distances, with phylogenetic trees generated via the *ape* package [45]. To test for overall differences in microbial community structure across treatments and time points, we applied permutational multivariate analysis of variance (PERMANOVA) using the **vegan** R package [46], applying Bonferroni correction to control for false positives. We further used PERMDISP2, a multivariate analogue of Levene’s test [47], to determine whether observed differences were influenced by dispersion within groups. Collectively, these analyses provided a robust framework for understanding the drivers of microbial community composition in our study.

To identify taxa associated with specific selection treatments or age groups, we performed Indicator Species Analysis using the **indicspecies** R package [48]. Based on the ASV table, we assessed associations between microbial taxa and either selection treatment or fly age. Indicator values were calculated using the `multipatt()` function, which evaluates both specificity and fidelity of taxa to groups, and statistical significance was assessed using 999 permutations.

To assess whether microbiome composition could predict fly age, dietary regime, or their interaction, we implemented a predictive random forest model adapted from [49]. Using the **caret** R package, we trained models to classify samples based on time point, treatment, and their combination, accounting for each unique vial and treatment-age group. Model performance was evaluated using repeated five-fold cross-validation, repeated five times. To account for variability in accuracy estimates, we ran 100 model iterations and calculated the mean predictive accuracy for each classification task. Additionally, we extracted the top 15 most important taxa contributing to each model, based on feature importance scores from the trained random forest classifiers.

## Results

### Overall longevity following diet selection

Lifespan performance in the non-selected base population (C) served as a benchmark for evaluating selection treatments (Fig. 2). Summary survival patterns are presented in (Additional file 2, Table S1). Overall, a log-rank test comparing survival across selection regimes revealed highly significant differences in lifespan (χ² = 274, df = 3, *p* < 2e–16). CH (i.e. constant high sugar) flies showed markedly lower survival (Fig. 2a-c), with far more deaths than expected under the null hypothesis of equal survival curves (median 27 days vs 42 days in C; Additional File 1, Table S2). In contrast, DA and FA (i.e. deteriorating- and fluctuating-availability) flies exhibited higher-than-expected survival (median 41 and 37 days, respectively). The control group lived the longest, significantly exceeding both CH (p = 5.91e-09) and FA (p = 0.0273) regimes (Fig. 2a). These patterns are reflected when survival is analyzed within 10-day bins around microbiome sampling time points, with the CH showing steeper mortality (Fig. 2c).

**Figure 2:**
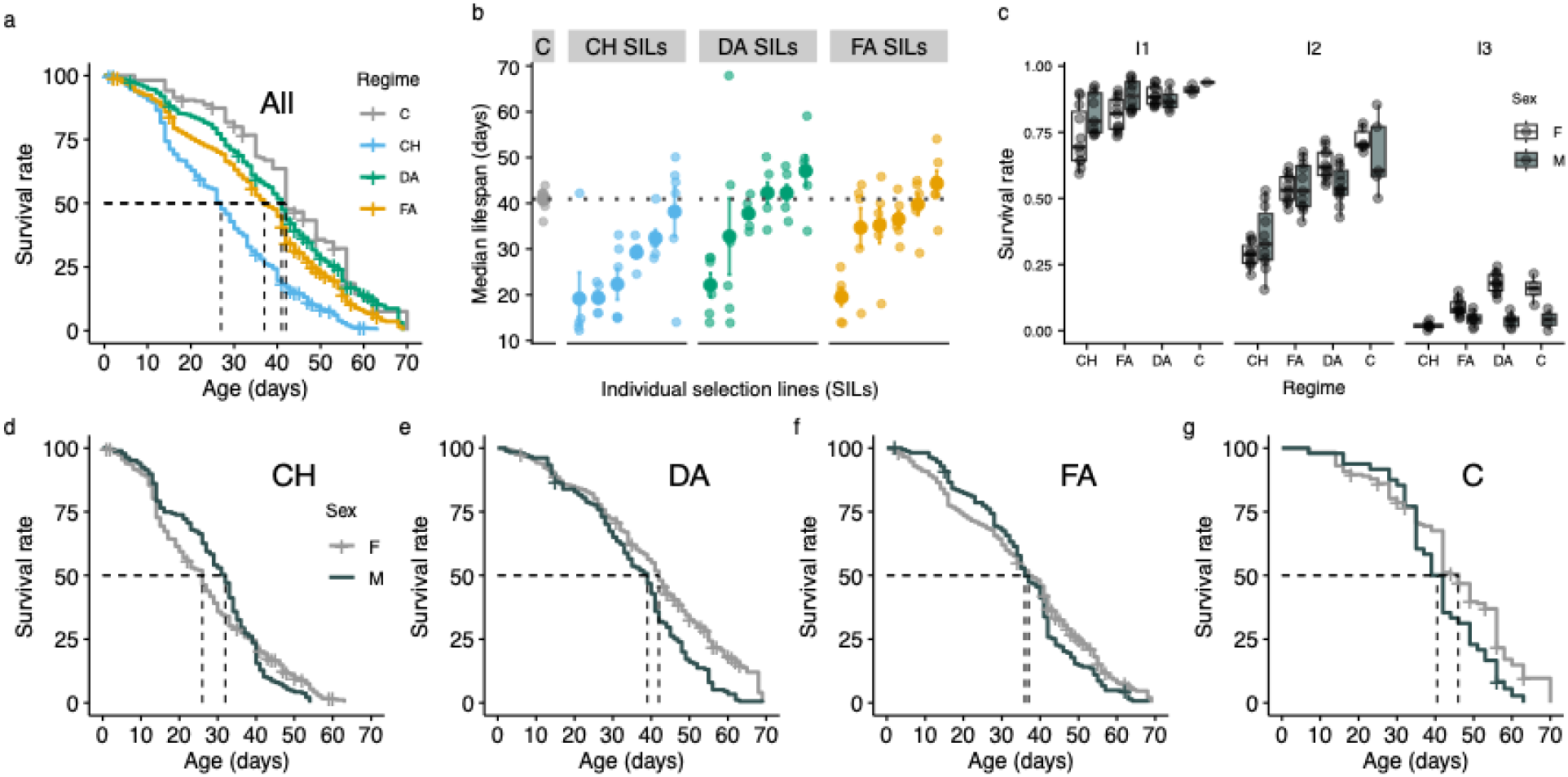
Survival performance: **a.** overall with both sexes together, **b.** median survival in each selection group of SILs, **c.** survival by sex based on bins of 10 days in which the sampling time point (of the microbiome) is the median (only bins containing assayed time points of 15, 35 and 60 days are shown (I1=10-20 days, I2=30-40 days, I3=55-65 days); **d-g**, comparison of sex performance averaged across SILs in each selection treatment.

### Sex-specific survival patterns

Sex differences in lifespan were apparent across all selection regimes. A Cox proportional hazards model with a time-varying effect of sex revealed significant differences in survival across regimes (*χ²* = 304.8, *df* = 4, *p* < 2e-16). Flies from the constant high-sugar regime (CH) had the highest risk of death (HR = 2.68, 95% CI: 2.28–3.15, *p* < 2e-16), followed by the fluctuating-availability (FA) regime (HR = 1.49, 95% CI: 1.27–1.75, *p* = 9.9e-07). The deteriorating-availability (DA) regime showed a modest, borderline effect (HR = 1.17, 95% CI: 1.00–1.38, *p* = 0.055). The time-varying term for sex, *tt(Sex_binary)*, was also significant (HR = 1.10, 95% CI: 1.06–1.13, *p* = 2.9e-10), indicating that the sex effect on mortality increased with age (Additional File 3 Fig. S1c). Males thus faced progressively higher mortality risk than females over time. Together, these results demonstrate that long-term selection under stressful dietary regimes, particularly constant high sugar, led to pronounced lifespan reductions, with sex-specific differences in mortality risk that intensified with age.

Although sex differences were clear, their magnitude and direction varied (Fig. 2d-g). In the control population, females outlived males, with median lifespans of 46 and 40.5 days, respectively (95% CI for females: 42 - 49; males: 35 - 44). A similar, though smaller, sex gap was observed under the DA regime (median: females 42, males 39 days). In the FA regime, survival differences between sexes were minimal, with overlapping confidence intervals and near-identical medians (females: 37 days, males: 36 days). Interestingly, under the CH regime, males lived slightly longer than females, with median lifespans of 32 vs 26 days. Generally, CH and FA treatments favored male survival earlier, reversing later in life with higher female survival (Fig. 2c, d, f). Conversely, DA and C regimes showed greater female survival through most of life (Fig. 2c, Fig. 2e,g, Additional file 3 Fig. S1 - S2), suggesting adjustment to dietary stress through reduced early mortality (DA, FA) or combined early and late-life mortality reduction (DA, C). These results indicate that selection regime influences not only overall lifespan but also modulates sex-specific survival patterns, with a marked sex-based lifespan reversal occurring under constant high sugar (CH). Results further suggest that dietary stress adaptation might involve distinct survival strategies in this population, with DA and FA regimes prioritizing early-life resilience and DA further extending late-life survival.

### Interactive impacts of sex and diet regime

Relative to C flies, the DA and FA regimes showed reduced early mortality (in I1 and I2, Fig. 2c) favoring longevity, but diverged later (in I3), with FA experiencing slightly higher late mortality while DA maintained slower rates. Control flies exhibited the lowest mortality in the first half of lifespan (I1 and I2). In contrast, CH flies suffered heavy mortality across the entire curve.

Underlying these differences are apparent rank changes in female vs male curves suggesting strong regime by sex interactions in survival (Fig. 2d-g). To evaluate the effects of selection regime, sex, and their interaction on mortality risk over time, we fit both Bayesian and frequentist generalized linear mixed models (GLMMs) using a complementary log-log link.

Estimates from both approaches were highly consistent, showing strong effects of time and the CH selection regime on increased mortality probability, while the DA and FA regimes had weaker or non-significant effects (Table 1, Fig. 3, Additional file 3 Fig. S3-S5). Male flies experienced a modestly elevated risk overall, and these differences were most pronounced in the CH treatment (Fig. 3). Bayesian posterior direction (PD) values further supported the robustness of key effects (e.g., time, CH, FA, Male, and CH: Male), with >96% of posterior samples in the direction of the mean. Full model estimates are provided in Table 1.

**Figure 3:**
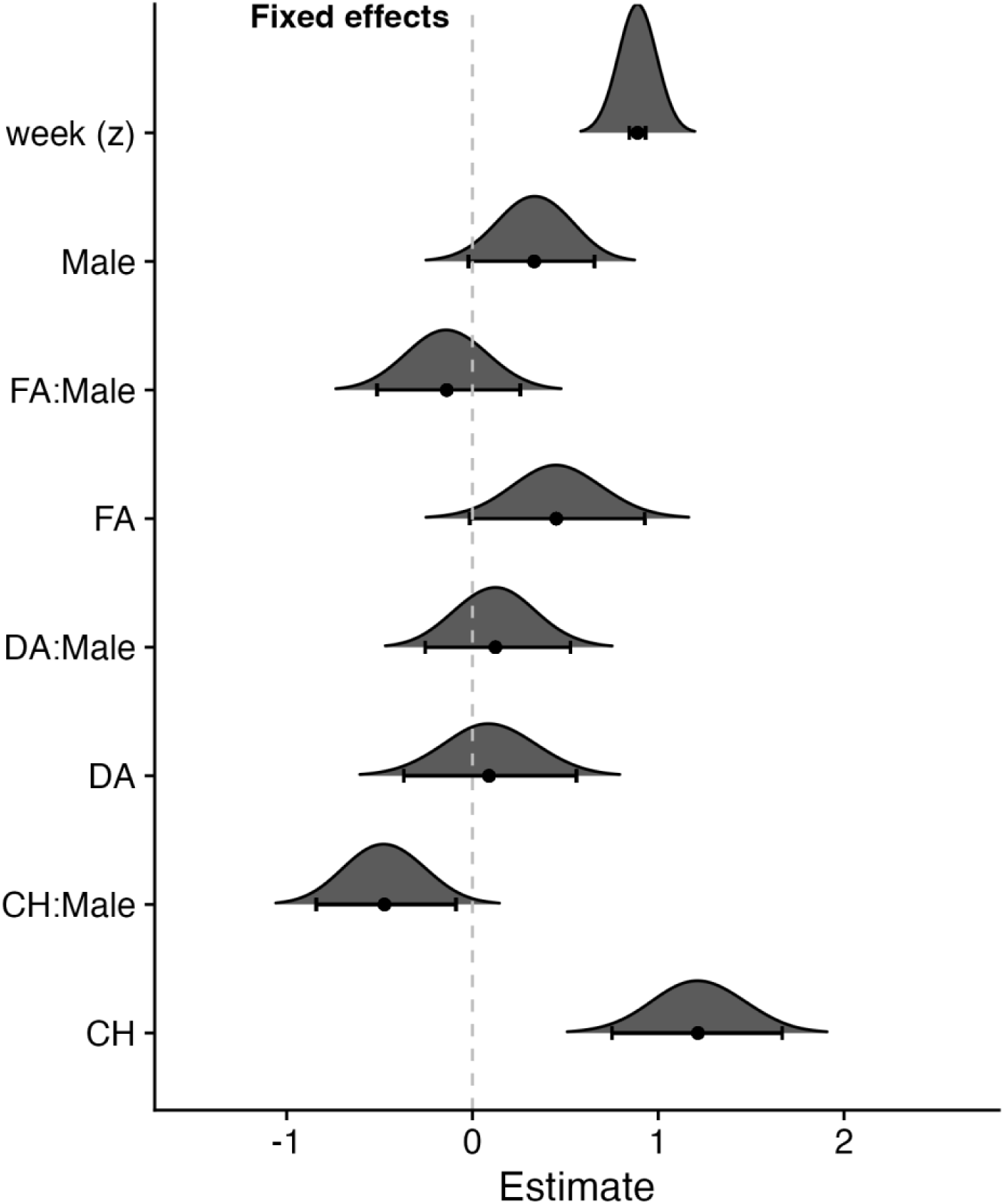
Fixed effects from hierarchical modeling of survival. Posterior distributions and 95% credible intervals (CIs) for fixed effects from the Bayesian model. Each ridge represents the estimated posterior distribution of a model coefficient. Points indicate posterior medians, and horizontal bars show 95% CIs. Vertical dashed line denotes the null effect (0). Parameters include treatment effects (CH, DA, FA), sex (Male), and their interactions, relative to the reference group (females in the control C regime). Estimates are on the scale of the linear predictor.

**Table 1.** Comparison of Bayesian and frequentist estimates from generalized linear mixed models (GLMMs) assessing the effects of selection regime, sex, and time (standardized week) on the probability of mortality. Bayesian estimates are posterior means with 95% credible intervals and posterior direction (PD), while frequentist GLMM estimates are shown with standard errors (SE), z-values, and associated p-values. The reference levels are the base population (C) and females.

| Effect | Bayesian estimate<br>(95% CI) | PD direction<br>(%) | glmer<br>estimate (se) | z-value | p-value |
| --- | --- | --- | --- | --- | --- |
| Intercept | -4.53 (-4.83, -4.23) | 100 | -4.54 (0.14) | -32.23 | < 2e-16 |
| Week (z) | 0.89 (0.85, 0.93) | 100 | 0.89 (0.02) | 40.39 | **< 2e-16** |
| CH | 1.21 (0.75, 1.67) | 100 | 1.23 (0.21) | 5.82 | **5.9e-09** |
| DA | 0.09 (-0.37, 0.56) | 65.4 | 0.11 (0.21) | 0.51 | 0.6084 |
| FA | 0.45 (-0.01, 0.93) | 97.2 | 0.47 (0.21) | 2.21 | **0.0273** |
| Male | 0.33 (-0.01, 0.67) | 96.7 | 0.35 (0.17) | 2.04 | **0.0416** |
| CH : Male | -0.47 (-0.84, -0.09) | 99.0 | -0.49 (0.19) | -2.55 | **0.0109** |
| DA : Male | 0.12 (-0.25, 0.53) | 73.4 | 0.10 (0.20) | 0.52 | 0.6007 |
| FA : Male | -0.14 (-0.51, 0.26) | 75.4 | -0.16 (0.20) | -0.81 | 0.4210 |

Predicted probabilities of death varied by selection regime and sex in both Bayesian and frequentist GLMMs (Additional File 3 Fig. S3, S4, S6). In C flies, females showed the lowest mortality probability (∼1%), while CH females had the highest (∼3–4%), with overlapping estimates between models. Males consistently exhibited higher predicted risks than females within most regimes, particularly under DA and FA. Both models revealed minimal sex differences in CH, where female and male predictions were comparably elevated. These adjusted predictions support the significant regime and sex effects identified in the full models.

### Microbial composition across selection regimes and time points

To assess microbial composition, we first analyzed the average relative abundance of phyla across time points and selection regimes (Fig. 4). Overall, all selection regimes (CH, DA, FA) displayed broadly similar microbial compositions that were distinct from the control, primarily due to lower Proteobacteria abundance and higher Firmicutes at the initial time point. Over time, Proteobacteria steadily increased across all treatments, ultimately converging between 45–65 % by T2. Other phyla, including Actinobacteria, Bacteroidota, and Fusobacteriota, were more abundant in the selected regimes than in controls but generally declined with age (Additional file 3: Fig. S8c–e).

**Figure 4:**
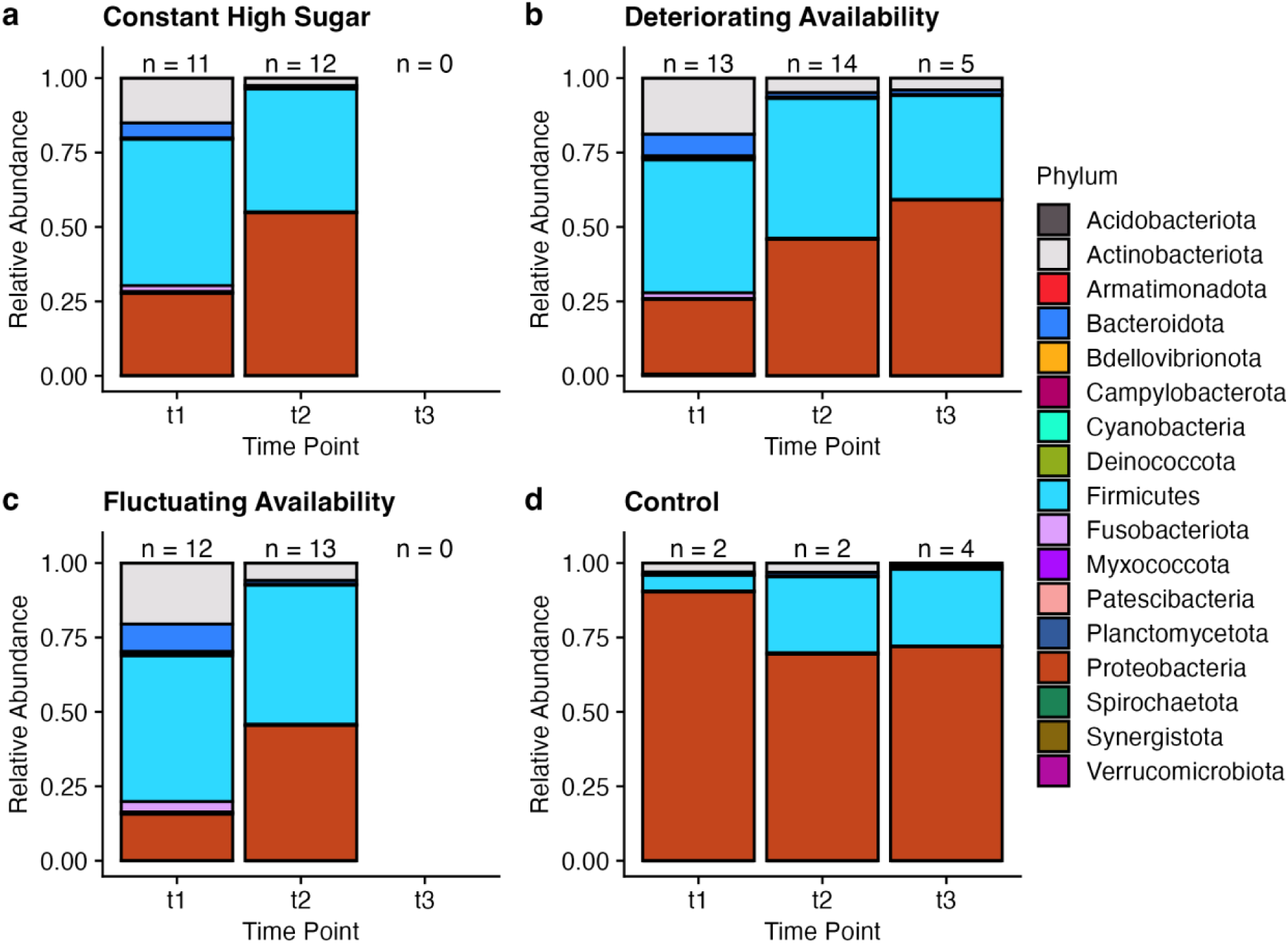
Phylum-level relative abundance of bacterial communities across timepoints and dietary treatments. Stacked bar plots show the log-transformed relative abundance of bacterial phyla at three timepoints t1, t2 and t3 (i.e. 15, 35, and 60 days post-eclosion) within each dietary treatment: **a.** constant high sugar (CH), **b.** decreasing nutrient availability (DA), **c.** fluctuating nutrient availability (FA), and **d.** control non-selected group (C). Each bar represents the mean relative abundance of samples collected at a given timepoint (n indicated above each bar). Colors correspond to bacterial phyla, as shown in the shared legend.

To explore compositional significance of observed relative abundance dynamics, we performed an indicator species analysis (ISA) at both the genus and species level. At the species level, an indicator species analysis (ISA) identified 35 species differing across time points, 5 across selection regimes, and 18 across both factors. Most time-related species (81 %) were enriched at the earliest time point (T1), highlighting pronounced early-life microbiome restructuring.

Control samples were enriched in three *Acetobacter* species (*A. aceti*, *A. oryzifermentans*, and *A. tropicalis*), whereas *Corynebacterium amycolatum* and *Mobiluncus curtisii* (Actinobacteria) were associated with CH and FA, respectively (Table 2). Several species were linked to specific combinations of diet and time. Notably, the genus of core Drosophila gut colonizers, *Acetobacter,* were depleted in the selected lines but present across all time points in controls, indicating that long-term dietary selection disrupted the *Acetobacter*-dominated microbiome characteristic of non-selected flies. The full output of the ISA looking across time point and selection regime can be found through (Additional file 4).

**Table 2:**
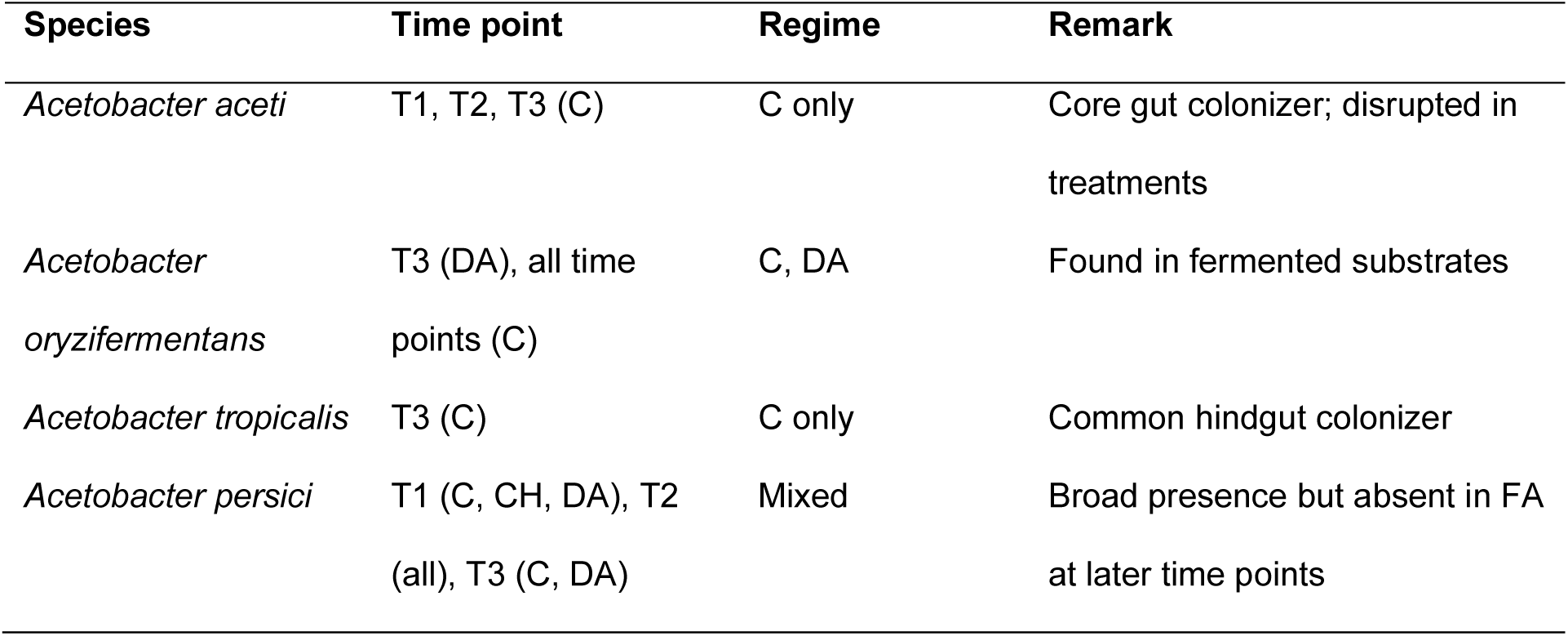
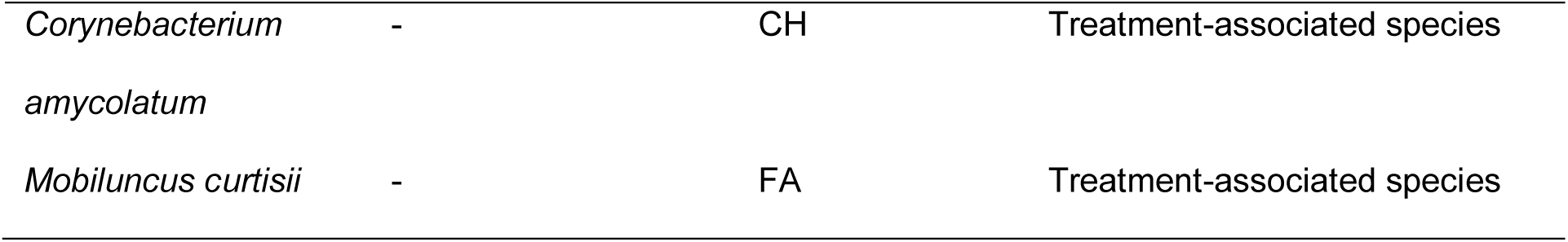
Species-level enrichment across time and selection regime.

At the genus level, ISA identified 39 genera differing across time points, 8 across selection regimes, and 28 across both factors (Additional file 2 Table S2). Among the diet-associated genera, *Fructilactobacillus* (Firmicutes) was enriched in CH; *Mobiluncus* (Actinobacteria) and *Pelomonas* (Proteobacteria) in FA; *Leptotrichia* (Proteobacteria) in CH + FA; *Acinetobacter* (Proteobacteria) in DA + FA; and *Actinomyces* (Actinobacteria), *Neisseria* (Proteobacteria), and *Lacticaseibacillus* (Firmicutes) in CH + DA + FA. Together, these results indicate that dietary selection history reshaped both phylum- and genus-level composition, resulting in distinct, convergent microbiomes among selected lines that diverged from the control.

Based on prior studies linking microbial composition to immune function [50, 51], we hypothesized that Proteobacteria and Firmicutes could influence hosts positively or negatively depending on their relative proportions. We calculated phyla ratios as age-specific indicators of treatment effects (Additional file 3 Fig. S8f-g). In the selection regimes, the Proteobacteria-Firmicute ratio generally increased over time (Additional file 3 Fig. S8f), while the Bacteriodota-Firmicute ratio decreased between T1 and T2 but then increased in DA between T2 to T3 (Additional file 3 Fig. S8g). The control group began with higher ratios than the treatments and exhibited a general decline, suggesting an age-related increase in Firmicutes relative to Proteobacteria and Bacteriodota. These patterns reveal contrasting dynamics between Proteobacteria and Bacteroidota that are influenced by Firmicutes.

## Microbial diversity dynamics and predictive modeling

We assessed microbial diversity across time points and dietary selection treatments using both alpha and beta diversity metrics. At the ASV level, both observed richness (Fig. 5a) and Shannon index (Fig. 5b), shifted significantly between T1 and T2 in all selection regimes, particularly in DA and FA indicating substantial early-life microbial restructuring. Some differences persisted into T3 in DA, although CH and FA lacked T3 data due to shortened lifespans. Similar patterns were observed at the genus- evel (Fig. 5c-d). Beta-diversity analyses (PCoA and PERMANOVA) revealed significant effects of time point, selection regime, and their interaction (p = 0.001), with pairwise contrasts showing pronounced T1 to T2 shifts in DA and FA (Fig. 6b-c). Dispersion tests further supported temporal changes within treatments, especially in DA and CH, and early compositional divergence between control and selected lines. Overall, age exerted a stronger influence than dietary regime on microbial community structure.

**Figure 5:**
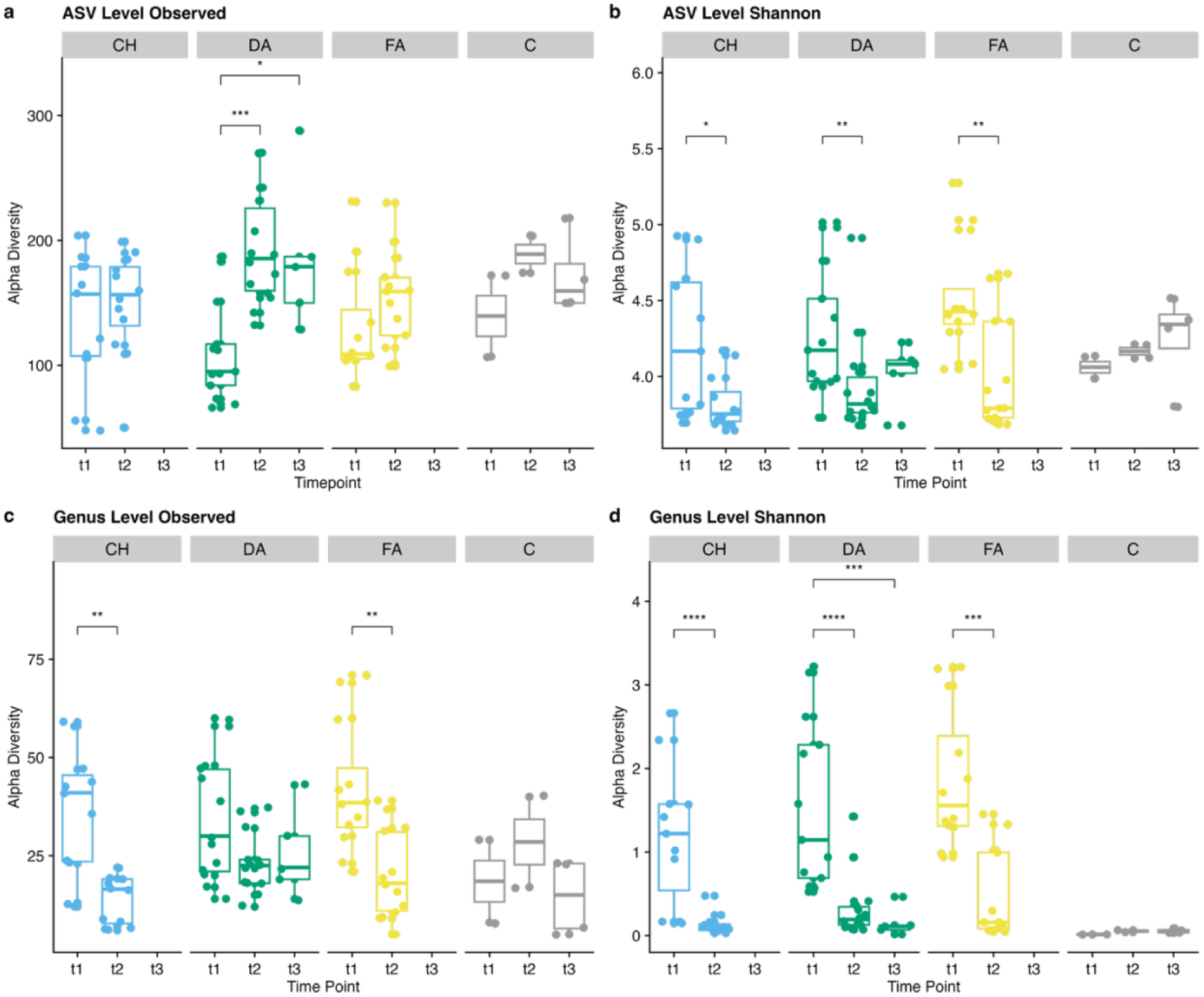
Temporal changes in microbial alpha diversity across dietary treatments(CH, DA, FA and C) at ASV (a-b) and genus (c-d) levels. **a&c** are observed diversity while **b&d** are Shannon-Weiner index. Significance between timepoints is from pairwise Wilcoxon rank-sum tests (*p* < 0.05), using raw p-values. Timepoints t1, t2 and t3 are 15, 35, and 60 days post-eclosion.

**Figure 6:**
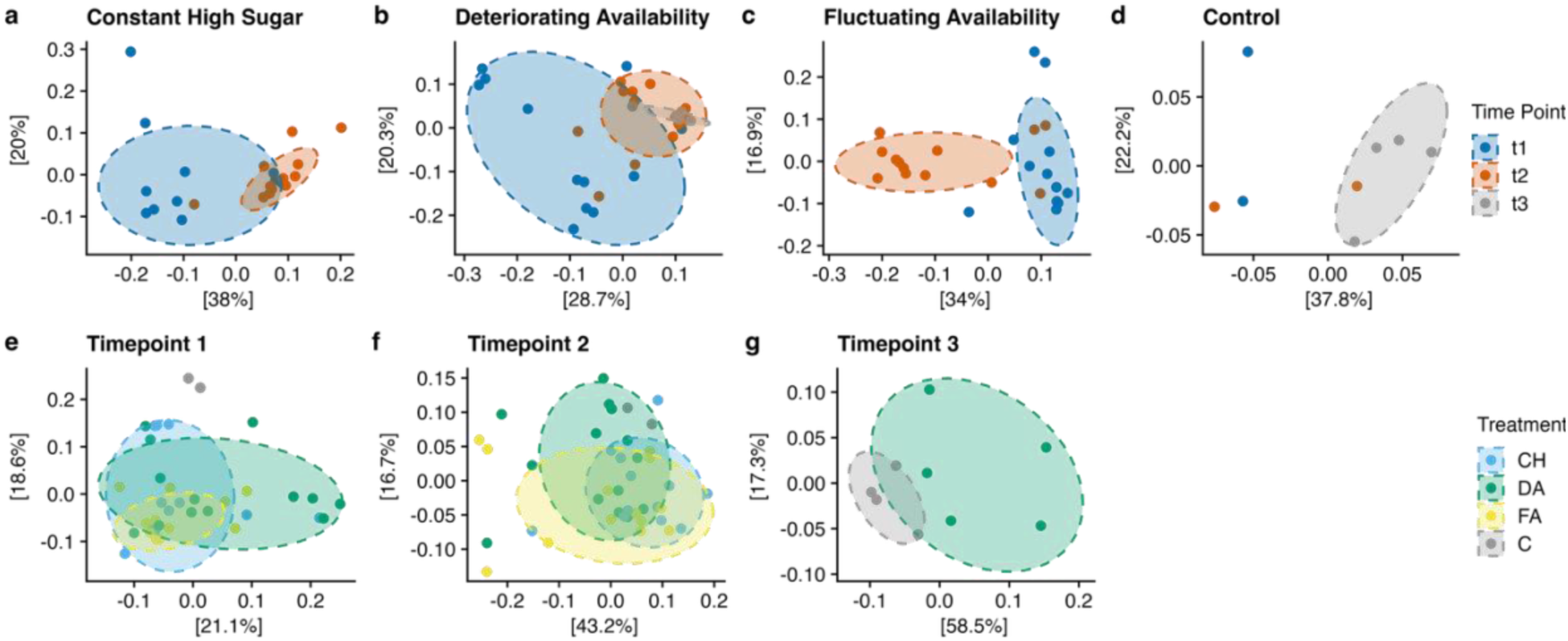
PCoA showing species diversity of dietary regimes over time (a - d) and across regimes at each timepoint (e - g). **a.** constant high sugar (CH), **b.** decreasing nutrient availability (DA), **c.** fluctuating nutrient availability (FA), **d.** control non-selected group (C), **e.** 15 days post eclosion (dpe) (T1), **f.** 35 dpe (T2), and **g.** 60 dpe (T3). CH and FA groups did not reach t3, hence no comparisons for T3 for a, c, and g. UniFrac distances are used as beta-diversity metric.

To evaluate the predictive value of microbial profiles, we trained random forest models to classify age, selection regime, and their interaction (Additional file 3 Fig. S10). The model predicting time point achieved the highest accuracy (78.8 %), identifying *Acetobacter persici*, *Cutibacterium acnes*, and *A. oryzifermentans* as key predictors. Models predicting dietary regime or the time x regime interaction performed less accurately (40.1 % and 32.2 %, respectively), likely reflecting limited sample sizes in the control and T3. Across all models, *A. persici* consistently emerged as a strong age-associated taxon, increasing in abundance in older flies across regimes.

Given its prominence in both random forest and indicator species analyses, we examined *Acetobacter* abundance in greater detail (Fig. 7). *A. persici* was the most abundant species across regimes, followed by *A. aceti* and *A. oryzifermentans*. From T1 to T2, *A. persici* increased significantly across all selected regimes but not in controls, while *A. aceti* rose in DA and FA. Between treatments, *A. aceti* differed significantly from controls in both T1 and T2, and *A. oryzifermentans* showed broader regime-specific differences. Together, these results indicate that age-related microbial shifts, dominated by *Acetobacter* taxa, are more consistent and predictable than those driven by dietary selection history. However, persistent differences in these core taxa between control and selected regimes suggest that long-term dietary selection also left distinct microbial legacies shaping community composition across ages.

**Figure 7:**
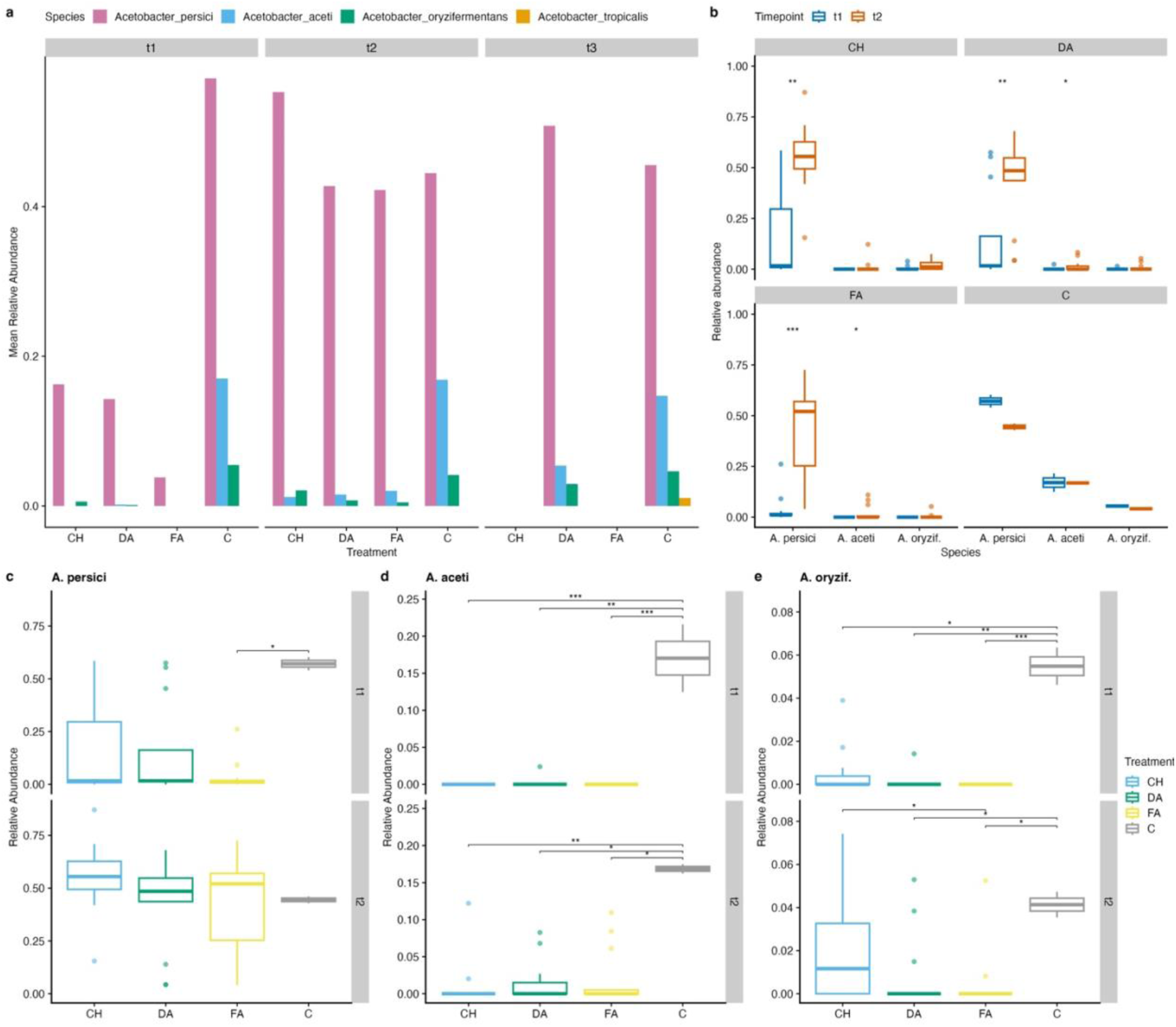
Prevalence of various Acetobacter species across samples: **a.** overview of the average relative abundance of four different acetobacter species. **b.** change in abundance of each species between 15 dpe (T1) and 35 dpe (T2) of each dietary regime. **c-e**. differences between dietary regimes in **c.** *A. aceti*, **d.** *A. persici*, and **e.** *A. oryzifermentans* at both T1 and T2. **b-e**. p-values from Wilcoxon ranked-sum tests.

## Discussion

We investigated how long-term dietary selection and inbreeding histories affect survival and microbial community dynamics in *Drosophila melanogaster*. To do so, we used individual diets we have characterized earlier in the DSPR (i.e. high sugar, reduced yeast, and standard diet) [31, 52, 53] to construct selection conditions, DA, FA and CH. While none of the selected lines outperformed the control in lifespan, DA (decreasing protein) selection, representing repeated exposure to a high-to-low dietary environment, produced lifespan performance comparable to the control group after 65 generations. In contrast, flies from FA and CH selection regimes exhibited reduced lifespan when maintained on the standard diet. This pattern indicates that chronic exposure to specific dietary stressors can leave lasting physiological and microbial legacies on lifespan, even after long-term rearing in a common environment. These findings extend our prior work and highlight that historical nutritional stress can generate enduring, and sometimes maladaptive, consequences for longevity.

## Longevity and sex-specific responses to dietary selection regimes

Among selection regimes, the constant high-sugar (CH) lines showed the strongest survival cost, particularly in females. Rather than indicating incomplete adaptation, this likely reflects a persistent metabolic legacy of chronic sugar exposure, which may influence host–microbe interactions. Prolonged selection under carbohydrate-rich conditions may have reinforced physiological priorities toward nutrient storage and reproduction at the expense of somatic maintenance, leading to reduced lifespan when flies are maintained on a more balanced diet [54, 55]. The pronounced early mortality in CH females suggests carryover effects of metabolic dysregulation during peak reproductive activity, consistent with sugar-induced insulin resistance observed in experimental evolution studies (e.g., [56]).

By contrast, lines exposed to fluctuating or deteriorating availability (FA and DA) exhibited more moderate declines, potentially reflecting historical exposure that promoted metabolic flexibility or resilience to nutrient variation rather than chronic abundance. In these lines, survival during early adulthood (a key reproductive window) was maintained, whereas later mortality increased, suggesting trade-offs in resource allocation between reproduction and maintenance.

Notably, sex-specific patterns emerged: FA males lived longer than females early in life, possibly facilitating sustained mating at a cost to female survival, while in DA a mildly opposite pattern was observed, consistent with maternal investment being favored under limited protein supply. Given that our data focus on longevity patterns rather than a complete set of life-history traits, we interpret these findings as reflecting evolved differences in physiological maintenance rather than comprehensive life-history dynamics. Further, although mutation accumulation is a theoretical factor influencing survival differences, it is unlikely to explain our results. The 65 generations of directional selection in large populations would have efficiently purged deleterious alleles. Thus, the observed survival differences more plausibly reflect evolved physiological adjustments to historical nutritional regimes rather than stochastic accumulation of mutations. Overall, the results point to divergent physiological trajectories rooted in past dietary environments, with lasting effects on survival even under a standardized diet.

### Microbiome responses to selection

Selection regimes also produced distinct microbiome profiles, representing the primary and most robust evidence of persistent effects of dietary selection. Compared to the control group, selected lines exhibited increased microbial richness and altered taxonomic composition, particularly early in life. The control flies maintained lower Firmicutes and higher Proteobacteria in early adulthood, consistent with a more beneficial, stable microbiome previously associated with greater lifespan in *Drosophila* [57, 58]. Selected lines experienced early colonization by taxa such as *Corynebacterium* and *Mobiluncus*, which in other systems include opportunistic or inflammation-associated species [16, 59, 60]. Studies in both flies and mammals show that early microbiota disruptions can impair host immunity and longevity [12]. These findings suggest that historical dietary stress can imprint persistent effects on microbiome assembly, potentially compromising host survival persistently after the selective pressure is withdrawn. The DA (and to a lesser degree, FA) lines, which maintained more survivors into later life, also showed similar dynamics in microbial composition across age. This is consistent with work showing that early adult microbiome establishment is critical for later-life health in flies [59, 61]. The ability to maintain beneficial taxa may therefore represent a retained trait from prior selection rather than ongoing adaptation in stressful nutritional environments.

We also detected consistent reductions in beneficial microbial taxa between the control and selection treatments in both T1 and T2, particularly in two core *Acetobacter* species: *A. aceti* and *A. oryzifermentans*. As members of the acetic acid bacteria group, *Acetobacter* species are integral to the fruit fly microbiome and have been linked to critical host functions, including growth and development, insulin signaling, lipid metabolism, and stem cell activity [62, 63]. In our study, flies from the selection regimes exhibited lower relative abundances of these microbes, suggesting that nutritional stress and inbreeding histories may have disrupted the core microbiota. Such disruptions could impair physiological processes and contribute to microbial imbalance and reduced survival. Interestingly, although *A. aceti* has been previously associated with shorter lifespan in *D. melanogaster* [64], we observed the highest levels of *A. aceti* and the longest lifespan in control flies. This contrast may reflect context-dependent consequences rather than directional adaptation, underscoring how dietary stress can interfere with host–microbiome relationships in ways that persist beyond the period of exposure.

Although flies were maintained for multiple generations on a standardized diet after the selection phase, microbiome differences among lines remained evident, suggesting that evolutionary history of selection constrained microbial assembly long after the environmental stress was removed.

### Patterns in dominant bacterial phyla

Patterns in dominant bacterial phyla further underscore the link between microbiome dynamics and host physiology. The ratio of Bacteroidota to Firmicutes decreased with age in all lines, probably suggesting an age-related shift in immune potential. Previous work associates higher Bacteroidota with immune activation [50] and lower Firmicutes with gut homeostasis [58]. The control flies, which had higher early Proteobacteria and lower Firmicutes, may have benefited from this composition. Conversely, historically stressed lines with elevated Firmicutes may carry immune and metabolic imbalances linked to prior selection, leading to greater community reorganization. We also observed an inverse relationship between Proteobacteria and Bacteroidota in selected lines, a pattern often linked to inflammation and metabolic stress [65, 66]. This microbial imbalance is well-documented in human and mouse studies under high-fat or sugar diets [67], suggesting deep evolutionary parallels in how dietary stress leaves microbial legacies across taxa.

### Microbiome succession and indicator taxa

Indicator species analysis showed rapid microbiome restructuring early in adulthood. Most microbial turnover occurred at T1, reinforcing the importance of early life as a window for microbiome establishment. While many groups declined over time, only a few were enriched in specific treatments, suggesting that historical exposure to dietary stress shaped which taxa persisted, rather than implying active adaptation of the microbial community. The presence of potentially pro-inflammatory species such as *Porphyromonas bennonis* and *Cutibacterium acnes* in some selection treatments raises further concern. These taxa are linked with chronic inflammation in mammals [68], and their persistence in flies may indicate long-term consequences for immune homeostasis. The apparent protective role of *Lactobacillus* in some groups aligns with studies highlighting its antagonistic effect on pathogens [69]. Thus, persistent survival differences among selection lines may result from retained microbial assemblages capable of modulating immune and metabolic balance, rather than ongoing adaptive co-evolution.

### Limitations and future directions

Our experimental design imposed a shared long-term post-selection environment, which could mask environment-specific adaptive responses. Moreover, fewer flies from the CH and FA groups survived to late age points, limiting effective comparisons at T3. Another limitation is that we did not characterize the initial microbiome prior to experimental evolution or immediately after the selection phase, which would have helped determine the magnitude of microbiome evolution in both the control and selected lines before rearing on NutriFly. Additionally, this experiment was not originally designed with microbiome evolution as a focal endpoint; thus, the lack of sterility during culture, the possible introduction of environmental microbes, and the use of food preservatives may have contributed to uncontrolled microbial variation. Nevertheless, detectable and consistent patterns across selection regimes and time points long after cessation of selection strengthen the inference that historical diet and subsequent inbreeding have left persistent legacies on both survival and microbiome structure.

Future work should disentangle host and microbial contributions to longevity by directly manipulating microbial communities in selection lines, ideally using gnotobiotic models. Reciprocal microbiome transplants between lines could further test whether microbial legacies from past selection can transfer phenotypes to naïve hosts. It would also be informative to re-examine the lines on their original selection diets and to assess whether the microbiota themselves evolved genetically or simply underwent species sorting. Finally, metagenomic profiling would clarify whether observed shifts reflect changes in microbial function or taxonomic identity.

## Conclusion

This study demonstrates that historical dietary selection and inbreeding leave persistent effects on the host-associated microbiome, which we consider the most robust outcome of this experiment. Selection lines exhibited microbial restructuring, and potential microbiome imbalance, while changes in survival and sex-specific life history strategies provide complementary context for interpreting these microbial patterns. These findings reinforce the idea that dietary history can imprint lasting host–microbiome legacies, influencing lifespan and physiological trajectories through both host genetic and microbial pathways. Given the rising relevance of diet-induced metabolic disorders in humans and pets, and changing climatic conditions for natural species, understanding how nutrition shapes microbial dynamics offers a promising path for translational insights.

## List of abbreviations

C: control group
CH: constant high availability
DA: deteriorating availability
FA: fluctuating availability
DSPR: Drosophila Synthetic Population Resource
SIL: selection inbred line
ΔAIC: delta Akaike Index
ASV: amplicon sequence variant
ISA: indicator species analysis

## Data availability

All 16S Illumina raw reeds are available in the NCBI Sequence Reads Archive (SRA) BioProject ID PRJNA1233815. Computer code to reproduce all analyses is available as a GitHub repository at https://github.com/nochet/selPhenotyping/ and analysis steps laid out in a ProjSteps.md file.

## Declarations

## Ethics approval and consent to participate

Not applicable. Clinical trial number: not applicable.

## Consent for publication

All authors give the publisher the right to publish this work.

## Acknowledgments

Authors thank Dr. Elizabeth G. King for discussions and Arthur K. Miller for technical assistance with experimental set up.

## Authors’ contributions

PKW performed 16S sequencing and analysis and wrote the manuscript. GM performed all fly experiments. GSK helped with data analysis and performed critical revision of the manuscript. EN conceived the study, designed experiments, analyzed data, and wrote the manuscript.

