## Additional File 1 for "Persistent effects of dietary selection and inbreeding on microbiome composition and longevity in Drosophila"

**GLMM RESULTS**

A. **Frequentist survival analysis: GLMM and Cox Frailty Model Comparison**

We analyzed survival in our experimental population using two complementary modeling approaches: a discrete-time generalized linear mixed survival model (GLMM) with a complementary log-log (cloglog) link and a mixed-effects Cox proportional hazards model (CoxME). Both models incorporated the fixed effects of regime, sex, and their interaction, with random intercepts for selection line to account for experimental replication (replicate nested in selection line, *sel_line*).

GLMM

1. Interaction model:

```
glm_interaction <- glmer(
  event ~ week_z + regime * Sex + (1 | sel_line),
  data = sel_long,
  family = binomial(link = "cloglog"),
  control = glmerControl(optimizer = "bobyqa", optCtrl = list(maxfun = 2e5)))
```

2. Nested model

```
glm_nested <- glmer(
  event ~ week_z + regime * Sex + (1 | sel_line/rept),
  data = sel_long,
  family = binomial(link = "cloglog"),
  control = glmerControl(optimizer = "bobyqa", optCtrl = list(maxfun = 2e5)))
```

Where time (week) is scaled:

week_z <- scale(week) # Mean 0, SD 1

3. Compare

anova(glm_interaction, glm_nested)

Adding the nested random effect ((1 | sel_line/rept)) does not improve model fit over just (1 | sel_line). The models have identical log-likelihoods, and:

- ΔAIC = +2 (a worse model)

- Chi-squared = 0, p = 1

- Either rept adds no additional variance in explaining mortality, or

- There's not enough replicate-level variation to estimate a second random effect reliably (especially if rept is not very replicated or contains sparse events per group).

Therefore, going forward, we removed nested replicates in all models. We compared a Cox and a GLMM without replicate nesting in random effect, selection line.

COX Frailty model

cox_frailty_me <- coxme(Surv(week, event) ~ regime * Sex + (1 | sel_line), data = sel_long)

#### Model Comparison (glmm vs coxme with ran effs but no nesting of reps)

The GLMM outperformed the CoxME model in terms of Akaike Information Criterion (AIC), with values of 18,794.6 and 41,313.7, respectively (Table 1). Cross-validation of test log-likelihoods further supported the superior predictive performance of the GLMM, with an average test log-likelihood of -1906.8 compared to -3475.5 for the CoxME (Table 1).

Table S1. Model fit statistics comparing GLMM and CoxME models for survival data.

| Model | AIC | Average Test Log-Likelihood (5-fold CV) |
| --- | --- | --- |
| GLMM (cloglog) | 18,794.6 | -1906.8 |
| CoxME (mixed effects) | 41,313.7 | -3475.5 |

### GLMM Results (cloglog link)

The best-fitting GLMM included week (z-scored), selection regime (CH, DA, FA vs. C), sex, and their interaction. Significant effects included time (week_z), the CH and FA regimes, male sex, and a negative interaction between the CH regime and male sex. The GLMM revealed significant main effects of regime and sex on survival probability (p < 0.05), and a significant regime × sex interaction (p < 0.01), indicating that the effect of selection regime on mortality risk differs between males and females (Table 2).

Table S2. Fixed effects estimates from GLMM for survival (cloglog link). Reference group: Regime_DA, Female.

| Effect | Estimate | Std. Error | z value | p-value |
| --- | --- | --- | --- | --- |
| (Intercept) | -4.54079 | 0.1409 | -32.227 | < 2e-16 |
| week_z | 0.88731 | 0.02197 | 40.391 | < 2e-16 |
| regimeCH | 1.23261 | 0.21181 | 5.819 | 5.91e-09 |
| regimeDA | 0.10925 | 0.21323 | 0.512 | 0.6084 |
| regimeFA | 0.46592 | 0.21103 | 2.208 | 0.0273 |
| SexM | 0.35316 | 0.17333 | 2.038 | 0.0416 |
| regimeCH:SexM | -0.49229 | 0.19341 | -2.545 | 0.0109 |
| regimeDA:SexM | 0.10251 | 0.19583 | 0.523 | 0.6007 |
| regimeFA:SexM | -0.15686 | 0.19495 | -0.805 | 0.4210 |

Model AIC: 18794.6; Log-likelihood: -9387.3

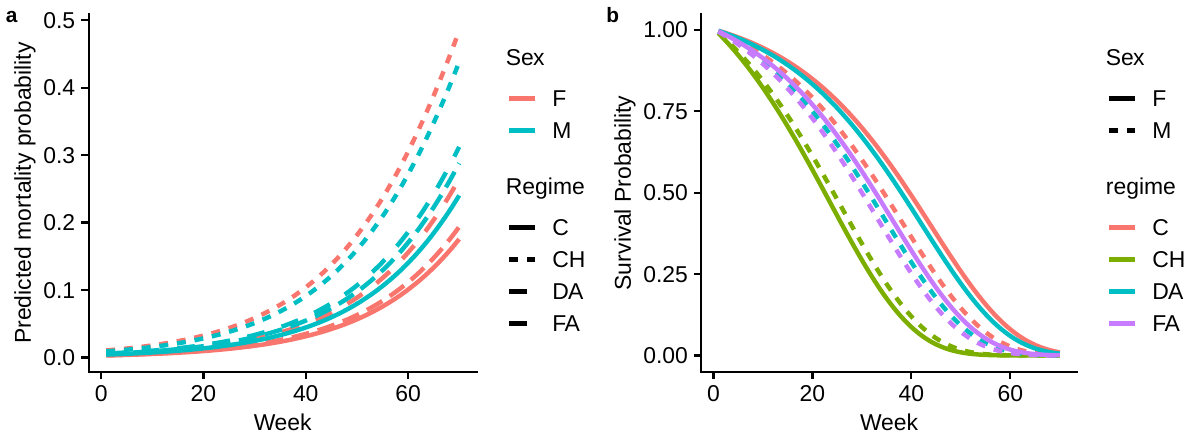

Predicted survival – coxme

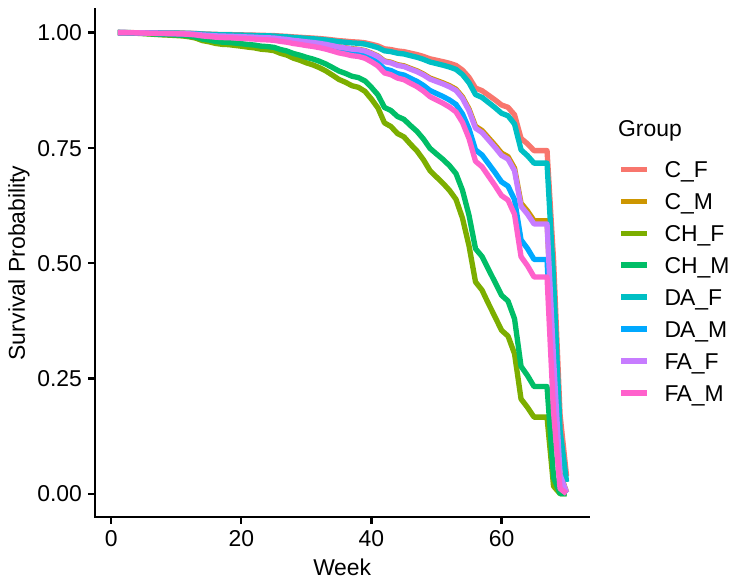

B. **Bayesian GLMMs with a cloglog link: Model specification, comparison and interpretation**

```
> summary(bayes_model)
 Family: bernoulli
  Links: mu = cloglog
Formula: event ~ week_z + regime * Sex + (1 | sel_line)
   Data: sel_long (Number of observations: 82731)
  Draws: 4 chains, each with iter = 4000; warmup = 1000; thin = 1;
         total post-warmup draws = 12000
```

Multilevel Hyperparameters:

```
~sel_line (Number of levels: 30)
              Estimate Est.Error l-95% CI u-95% CI Rhat Bulk_ESS Tail_ESS
sd(Intercept)     0.43      0.07     0.31     0.59 1.00     4082     6140
```

Regression Coefficients:

```
              Estimate Est.Error l-95% CI u-95% CI Rhat Bulk_ESS Tail_ESS
Intercept        -4.53      0.15    -4.83    -4.23 1.00     8118     8208
week_z            0.89      0.02     0.85     0.93 1.00    14793     8702
regimeCH          1.22      0.24     0.75     1.67 1.00     6556     7047
regimeDA          0.09      0.24    -0.37     0.56 1.00     6592     7344
regimeFA          0.46      0.23    -0.01     0.93 1.00     6411     6659
SexM              0.33      0.17    -0.02     0.66 1.00     7352     7207
regimeCH:SexM    -0.47      0.19    -0.84    -0.09 1.00     8013     7758
regimeDA:SexM     0.12      0.20    -0.25     0.53 1.00     7792     8315
regimeFA:SexM    -0.14      0.20    -0.51     0.26 1.00     7771     8322
```

Draws were sampled using sampling(NUTS). For each parameter, Bulk_ESS

and Tail_ESS are effective sample size measures, and Rhat is the potential

scale reduction factor on split chains (at convergence, Rhat = 1).

```
> summary(model_no_interaction)
 Family: bernoulli
  Links: mu = cloglog
Formula: event ~ week_z + regime + Sex + (1 | sel_line)
   Data: sel_long (Number of observations: 82731)
  Draws: 4 chains, each with iter = 4000; warmup = 1000; thin = 1;
         total post-warmup draws = 12000
```

Multilevel Hyperparameters:

```
~sel_line (Number of levels: 30)
              Estimate Est.Error l-95% CI u-95% CI Rhat Bulk_ESS Tail_ESS
sd(Intercept)     0.43      0.07     0.32     0.59 1.00     3326     5512
```

Regression Coefficients:

```
          Estimate Est.Error l-95% CI u-95% CI Rhat Bulk_ESS Tail_ESS
Intercept    -4.48      0.15    -4.78    -4.19 1.00     6477     7782
week_z        0.88      0.02     0.84     0.92 1.00    14884     9372
regimeCH      1.09      0.23     0.63     1.55 1.00     4953     5826
regimeDA      0.13      0.23    -0.32     0.59 1.00     5199     6585
regimeFA      0.42      0.22    -0.03     0.87 1.00     5289     6626
SexM          0.16      0.05     0.06     0.25 1.00    14043     8416
```

Draws were sampled using sampling(NUTS). For each parameter, Bulk_ESS

and Tail_ESS are effective sample size measures, and Rhat is the potential

scale reduction factor on split chains (at convergence, Rhat = 1).

```
> posterior_summary(model_no_interaction)
                                   Estimate  Est.Error          Q2.5         Q97.5
b_Intercept                   -4.482245e+00 0.14800606 -4.777983e+00 -4.192358e+00
b_week_z                       8.809061e-01 0.02172900  8.389747e-01  9.231465e-01
b_regimeCH                     1.089056e+00 0.23347979  6.343178e-01  1.550006e+00
b_regimeDA                     1.303180e-01 0.23032921 -3.228503e-01  5.878804e-01
b_regimeFA                     4.219565e-01 0.22476561 -2.619467e-02  8.705015e-01
b_SexM                         1.593899e-01 0.04923780  6.268457e-02  2.547483e-01
sd_sel_line__Intercept         4.272461e-01 0.06930405  3.150896e-01  5.871009e-01
Intercept                     -3.989945e+00 0.09582867 -4.179080e+00 -3.800860e+00
r_sel_line[1.DA,Intercept]    -6.876196e-02 0.19480911 -4.517043e-01  3.209440e-01
r_sel_line[10DA,Intercept]     5.821968e-01 0.19618033  1.973927e-01  9.774627e-01
r_sel_line[10FA,Intercept]     3.646681e-01 0.19118273 -1.848574e-02  7.459085e-01
r_sel_line[11.CH,Intercept]   -1.697487e-01 0.19292868 -5.543059e-01  2.074346e-01
r_sel_line[11FA,Intercept]    -8.402648e-02 0.18902020 -4.549072e-01  2.917290e-01
r_sel_line[12FA,Intercept]    -2.218415e-01 0.19061306 -5.945643e-01  1.485056e-01
r_sel_line[1FA,Intercept]      2.668215e-01 0.18828830 -1.024027e-01  6.417688e-01
r_sel_line[2.CH,Intercept]    -6.730466e-01 0.19287401 -1.055135e+00 -2.998678e-01
r_sel_line[2DA,Intercept]     -2.458902e-01 0.20074815 -6.412092e-01  1.486077e-01
r_sel_line[2FA,Intercept]      3.182440e-02 0.19051608 -3.433340e-01  4.042452e-01
r_sel_line[3CH,Intercept]     -1.591017e-01 0.19480695 -5.549687e-01  2.203430e-01
r_sel_line[4CH,Intercept]      5.306992e-01 0.19064295  1.569687e-01  9.050749e-01
r_sel_line[6DA,Intercept]      8.590895e-01 0.19631815  4.770877e-01  1.255617e+00
r_sel_line[7DA,Intercept]     -3.472573e-01 0.19914436 -7.396559e-01  4.482925e-02
r_sel_line[8CH,Intercept]      3.025691e-02 0.19338393 -3.548060e-01  4.123646e-01
r_sel_line[8FA,Intercept]     -3.636821e-01 0.18949651 -7.426009e-01  8.814037e-03
r_sel_line[9.CH,Intercept]     4.659737e-01 0.19245608  9.189057e-02  8.435043e-01
r_sel_line[9DA,Intercept]     -7.771311e-01 0.21642952 -1.214959e+00 -3.619051e-01
r_sel_line[sp1_44,Intercept]  -3.508131e-01 0.26305140 -8.855566e-01  1.466980e-01
r_sel_line[sp1_65,Intercept]  -6.435697e-02 0.23305752 -5.261721e-01  3.801211e-01
r_sel_line[sp2_37,Intercept]   1.677452e-01 0.27461569 -3.672809e-01  7.030433e-01
r_sel_line[sp2_66,Intercept]  -5.850028e-02 0.23322793 -5.272463e-01  3.887661e-01
r_sel_line[sp3_14,Intercept]  -8.846974e-03 0.23960466 -4.914065e-01  4.451649e-01
r_sel_line[sp3_20,Intercept]   6.063593e-02 0.23278033 -4.040215e-01  5.128521e-01
r_sel_line[sp4_2,Intercept]    5.097282e-02 0.26444993 -4.903339e-01  5.560162e-01
r_sel_line[sp4_53,Intercept]   2.429875e-01 0.26552221 -2.843063e-01  7.644384e-01
r_sel_line[sp5_114,Intercept] -3.218487e-01 0.25357972 -8.418216e-01  1.609821e-01
r_sel_line[sp5_86,Intercept]   1.731774e-01 0.24714990 -3.154800e-01  6.555407e-01
r_sel_line[sp6_106,Intercept]  3.727122e-02 0.21098036 -3.836560e-01  4.369798e-01
r_sel_line[sp6_93,Intercept]  -3.934405e-03 0.21944107 -4.351596e-01  4.201976e-01
lprior                        -1.274383e+01 0.08289998 -1.293057e+01 -1.260499e+01
lp__                          -9.423854e+03 5.37787728 -9.435323e+03 -9.414164e+03
z_1[1,1]                      -1.662179e-01 0.45601636 -1.051897e+00  7.359746e-01
z_1[1,2]                       1.392397e+00 0.49350612  4.301057e-01  2.370070e+00
z_1[1,3]                       8.734148e-01 0.46416028 -4.192682e-02  1.782147e+00
z_1[1,4]                      -4.059941e-01 0.45396081 -1.299914e+00  4.745122e-01
z_1[1,5]                      -2.004502e-01 0.44148022 -1.059593e+00  6.657582e-01
z_1[1,6]                      -5.302335e-01 0.44970381 -1.418535e+00  3.385951e-01
z_1[1,7]                       6.392964e-01 0.44824572 -2.304425e-01  1.520233e+00
z_1[1,8]                      -1.611228e+00 0.50467549 -2.619945e+00 -6.513401e-01
z_1[1,9]                      -5.895174e-01 0.47636977 -1.520934e+00  3.364867e-01
z_1[1,10]                      7.710227e-02 0.44593423 -8.032187e-01  9.508664e-01
z_1[1,11]                     -3.798598e-01 0.45737203 -1.293504e+00  5.085802e-01
z_1[1,12]                      1.271029e+00 0.48061434  3.460651e-01  2.235415e+00
z_1[1,13]                      2.054790e+00 0.53431850  1.040346e+00  3.126965e+00
z_1[1,14]                     -8.326246e-01 0.48137509 -1.767389e+00  9.987670e-02
z_1[1,15]                      7.256688e-02 0.45210762 -8.127407e-01  9.595808e-01
z_1[1,16]                     -8.699721e-01 0.45645849 -1.753836e+00  1.938474e-02
z_1[1,17]                      1.115603e+00 0.47434615  2.107537e-01  2.057947e+00
z_1[1,18]                     -1.855305e+00 0.55156520 -2.951148e+00 -8.031264e-01
z_1[1,19]                     -8.279029e-01 0.61858064 -2.083787e+00  3.555121e-01
z_1[1,20]                     -1.520005e-01 0.55334694 -1.247208e+00  9.008882e-01
z_1[1,21]                      3.944811e-01 0.64998295 -9.045583e-01  1.677336e+00
z_1[1,22]                     -1.379135e-01 0.55473550 -1.266318e+00  9.257083e-01
z_1[1,23]                     -2.080777e-02 0.56872203 -1.164297e+00  1.077587e+00
z_1[1,24]                      1.433323e-01 0.55275566 -9.506642e-01  1.234133e+00
z_1[1,25]                      1.228511e-01 0.63073663 -1.175274e+00  1.329681e+00
z_1[1,26]                      5.734709e-01 0.62810117 -6.773143e-01  1.790054e+00
z_1[1,27]                     -7.598510e-01 0.59683107 -1.975978e+00  3.851151e-01
z_1[1,28]                      4.103516e-01 0.58699867 -7.745672e-01  1.544830e+00
z_1[1,29]                      8.791255e-02 0.50137406 -9.198141e-01  1.043748e+00
z_1[1,30]                     -9.997472e-03 0.52190887 -1.036223e+00  9.961740e-01
```

> plot(model_no_interaction)

Hit <Return> to see next plot:

> pp_check(model_no_interaction, type = "stat", ndraws = 500)

> pp_check(model_no_interaction, type = "bars", ndraws = 200)

> model_comparison <- bayes_factor(bayes_model, model_no_interaction)

Iteration: 1

Iteration: 2

Iteration: 3

Iteration: 4

Iteration: 5

Iteration: 1

Iteration: 2

Iteration: 3

Iteration: 4

> model_comparison

Estimated Bayes factor in favor of bayes_model over model_no_interaction: 45.21841

This means that the full model (bayes_model), which includes the interaction between regime and Sex, is ~45 times more likely than the reduced model without that interaction.

According to common interpretative guidelines (e.g., Kass & Raftery, 1995):

| Bayes Factor (BF₁₀) | Strength of evidence for H₁ (full model) |
| --- | --- |
| 1 – 3 | Anecdotal |
| 3 – 10 | Moderate |
| 10 – 30 | Strong |
| 30 – 100 | **Very strong** |
| > 100 | Extreme |

Thus, a BF of 45.2 provides **very strong evidence** that including the regime:Sex interaction improves the model (i.e., this interaction explains important variation in event probability).

Our logistic model with a complementary log-log link suggests that the effect of selection regime on event probability differs by sex. This interaction is important enough to greatly improve predictive accuracy, as judged by Bayesian model comparison.
