## Additional File 2 for "Persistent effects of dietary selection and inbreeding on microbiome composition and longevity in Drosophila"

**Table S1: Survival rates in each selection group all flies and separate sex groups**

| Selection regime | Survival (%) | All  days (CI) | Female  days (CI) | Male  days (CI) |
| --- | --- | --- | --- | --- |
| CH | 90 | 11(9-12) | 10(8-12) | 12(10-14) |
|  | 75 | 14(14-15) | 14(14-15) | 16(14-23) |
|  | 50 | 27(26-28) | 26(23-27) | 32(29-33) |
|  | 25 | 39(36-40) | 39(36-40) | 39(36-40) |
|  | 10 | 49(47-52) | 50(48-54) | 43(41-48) |
| DA | 90 | 14(13-16) | 14(12-17) | 15(13-17) |
|  | 75 | 28(26-29) | 28(26-31) | 27(22-29) |
|  | 50 | 41(39-42) | 42(41-44) | 39(34-41) |
|  | 25 | 54(50-55) | 55(54-58) | 47(44-49) |
|  | 10 | 64(62-68) | 68(65-68) | 55(52-56) |
| FA | 90 | 12(11-14) | 11(8-13) | 16(14-17) |
|  | 75 | 21(18-25) | 19(16-23) | 27(23-31) |
|  | 50 | 37(35-40) | 37(35-40) | 36(35-40) |
|  | 25 | 49(47-51) | 50(48-53) | 44(42-48) |
|  | 10 | 58(56-60) | 58(56-62) | 55(51-57) |
| C | 90 | 23(16-28) | 18(14-28) | 28(16-32) |
|  | 75 | 35(30-35) | 35(28-39) | 35(32-37) |
|  | 50 | 42(42-49) | 46(42-49) | 40(35-44) |
|  | 25 | 56(56-56) | 56(56-58) | 49(42-56) |
|  | 10 | 63(60-NA) | 63(63-NA) | 56(56-NA) |

**Tables S2: Summaries of microbial diversity analyses**

### A) Alpha diversity (observed richness and Shannon index)

| Taxonomic level | Measure | Comparison | p-value | Remark |
| --- | --- | --- | --- | --- |
| ASV | Observed Richness | DA: T1 vs T2 | 0.0003 | Significant decline from T1 to T2 |
| ASV | Observed Richness | DA: T1 vs T3 | 0.02 | Significant decline |
| ASV | Shannon Index | CH: T1 vs T2 | 0.027 | Early-life shift in diversity |
| ASV | Shannon Index | DA: T1 vs T2 | 0.0091 |  |
| ASV | Shannon Index | FA: T1 vs T2 | 0.003 |  |
| Genus | Observed Richness | CH: T1 vs T2 | 0.0015 | Significant decline |
| Genus | Observed Richness | FA: T1 vs T2 | 0.0012 |  |
| Genus | Shannon Index | CH: T1 vs T2 | 6.7e-05 |  |
| Genus | Shannon Index | DA: T1 vs T2 | 2.7e-05 |  |
| Genus | Shannon Index | FA: T1 vs T2 | 0.00019 |  |
| Genus | Shannon Index | DA: T1 vs T3 | 0.00023 | Late-life change |

### B) Beta Diversity (PERMANOVA and Dispersion Tests)

| Comparison Type | Comparison | Test | p-value | Notes |
| --- | --- | --- | --- | --- |
| PERMANOVA | Selection regime | PERMANOVA | 0.001 | Significant |
| PERMANOVA | Time point | PERMANOVA | 0.001 | Significant |
| PERMANOVA | Interaction: Selection × Time | PERMANOVA | 0.001 | Significant |
| Pairwise | DA: T1 vs T2 | PERMANOVA | 0.045 | Significant within DA |
| Pairwise | FA: T1 vs T2 | PERMANOVA | 0.045 | Significant within FA |
| Dispersion | CH: T1 vs T2 | Permutation | 0.001 | Significant change in spread |
| Dispersion | DA: T1 vs T2 | Permutation | 0.001 |  |
| Dispersion | DA: T1 vs T3 | Permutation | 0.001 |  |
| Dispersion | FA: T1 vs T2 | Permutation | 0.003 |  |
| Dispersion | CH vs C at T1 | Permutation | 0.001 | Treatment-control difference |
| Dispersion | DA vs C at T1 | Permutation | 0.001 |  |
| Dispersion | FA vs C at T1 | Permutation | 0.001 |  |
| Dispersion | DA vs FA at T1 | Permutation | 0.034 |  |

### C) Prediction Accuracy of Random Forest Models

| Model Type | Prediction Target | Average Accuracy (%) | Key Predictive Species |
| --- | --- | --- | --- |
| Model 1 | Time Point | 78.8 | A. persici, Cutibacterium acnes, A. oryzifermentans |
| Model 2 | Dietary Selection Regime (CH, DA, FA) | 40.1 | A. persici, A. aceti, unidentified species |
| Model 3 | Time Point × Regime (T1, T2 only) | 32.2 | A. persici, unidentified species |
