## Additional File 3 for "Persistent effects of dietary selection and inbreeding on microbiome composition and longevity in Drosophila"

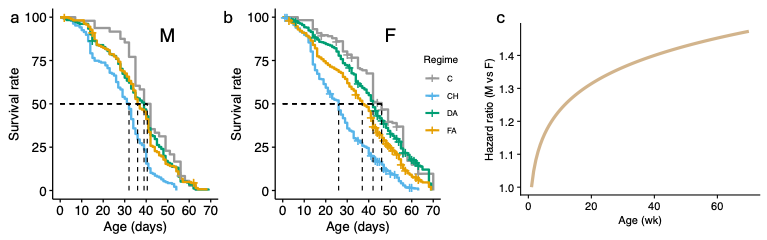

**Figure S1**: Survival patterns in long-term diet selected *Drosophila melanogaster*: **a.** Kaplan-Meir survival curves plotted by sex, **a.** males (M), **b.** female (F). Selection regimes: C, control (unselected); DA, deteriorating availability; FA, fluctuating availability; and CH, constant high sugar. **c.** Time-varying Cox model showed that sex effects on mortality increased with age (HR = 1.10, 95% CI: 1.06–1.13, p = 2.9e-10).

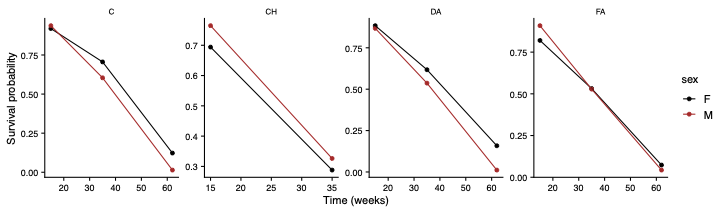

**Figure S2:** Survival patterns following selection in each regime visualized by sex focusing on time points evaluated in microbiome analysis. Notice changes in rank where C and DA females live longer vs CH which live shorter, and FA showing an interaction. Note two time points in CH as this group did not reach T3.

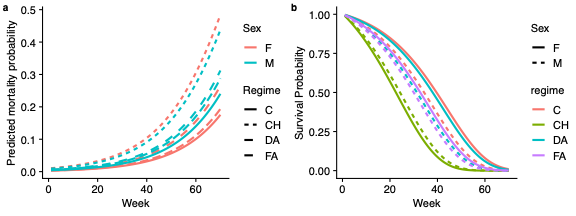

**Figure S3**: Predicted mortality and survival probabilities over time in each selection regime by sex obtained from a frequentist GLMM (see Methods for details).

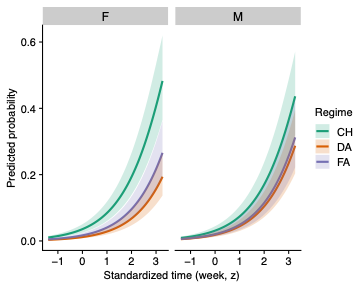

**Figure S4**: Predicted mortality probability over time in each selection regime by sex obtained from a corresponding Bayesian GLMM (see Methods for details).

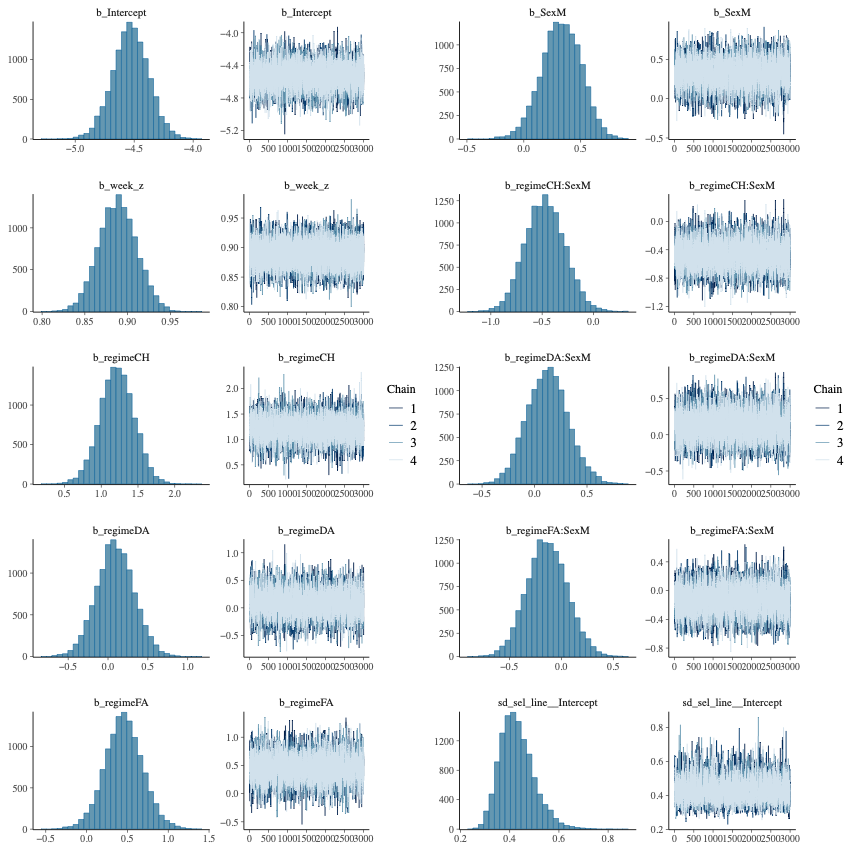

**Figure S5**: Trace plots and posterior density distributions for parameters estimated in the Bayesian mixed-effects cloglog model (event ~ week_z + regime * Sex + (1 | sel_line)) fit to the binary event data (n = 82,731). The plots show well-mixed chains across four sampling chains (12,000 post-warmup samples total), indicating good convergence and adequate sampling. All R̂ values were < 1.01, with no divergent transitions.

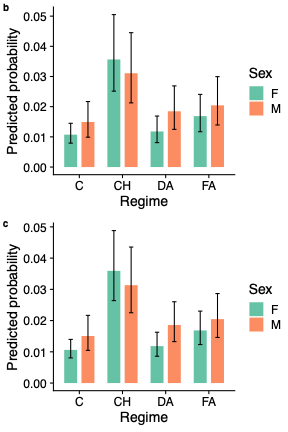

**Figure S6:** Predicted probabilities of mortality across selection regimes and sexes from Bayesian and frequentist generalized linear mixed models: **b.** Posterior estimates showing marginal effects of the interaction between selection regime and sex, **c.** Corresponding marginal means and confidence intervals from a glmer fit with the same fixed and random effects structure. Bars represent predicted probabilities. Error bars indicate 95% credible intervals (top) or 95% confidence intervals (bottom). Estimate look almost identical.

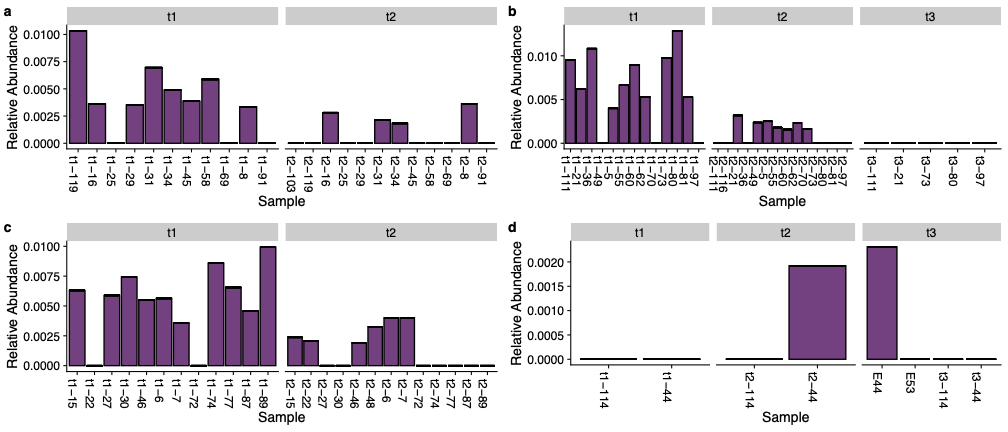

**Figure S7**: Genus-level log-transformed relative abundance of *Wolbachia*, showing the prevalence of *Wolbachia* across samples at both timepoints and dietary treatments. *Wolbachia* frequency is plotted by dietary treatment: a. constant high sugar (CH), b. decreasing nutrient availability (DA), c. fluctuating nutrient availability (FA), and d. control non-selected group (C).

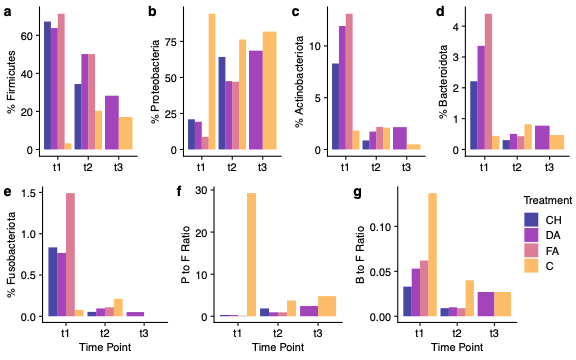

**Figure S8**: Changes in the percentage of phyla over time (t1-t3) among dietary regimes (a-e): **a.** Firmicutes, **b.** Proteobacteria, **c.** Actinobacteriota, **d.** Bacteroidota, and **e.** Fusobacteriota. Phyla ratios over time between dietary treatments (f-g): **f.** Proteobacteria to Firmicutes, **g.** Bacteroidota to Firmicutes. Time points: t1 - 15 days; t2 - 35 days; and t3 - 60 days.

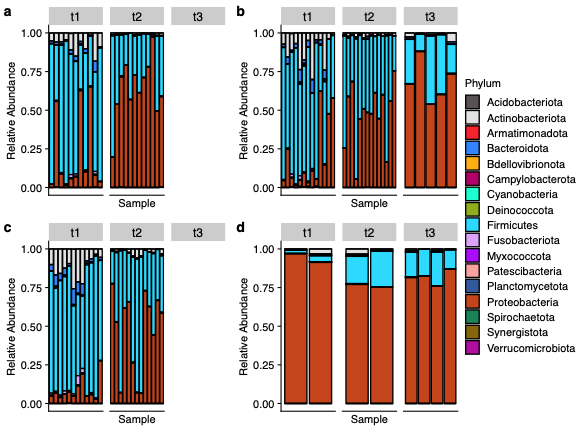

**Figure S9**: Relative abundance across all samples, separated by both timepoint and dietary treatment: a. constant high sugar (CH), b. decreasing nutrient availability (DA), c. fluctuating nutrient availability (FA), and d. control (C). Colors correspond to bacterial phyla, as shown in the shared legend. The absence of samples in a timepoint is represented by an empty plot in t3 of both CH and FA.

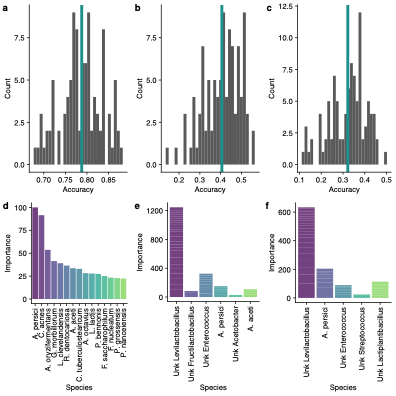

**Figure S10:** Random Forest predictive model average accuracy over one hundred predictions (a-c) and species identified that were important for predictive power in each model (d-f). Models include time point (a, d), selection regime (b, e), and time point and selection regime excluding T3 (c, f). *unk* denotes unidentified/unknown species of bacteria.
